# The primary donor of far-red Photosystem II: Chl_D1_ or P_D2_?

**DOI:** 10.1101/2020.04.03.021097

**Authors:** Martyna Judd, Jennifer Morton, Dennis Nürnberg, Andrea Fantuzzi, A. William Rutherford, Robin Purchase, Nicholas Cox, Elmars Krausz

**Affiliations:** Research School of Chemistry, Australian National University, Canberra 2600, Australia; Department of Life Sciences, Imperial College, London SW7 2AZ, United Kingdom

**Keywords:** photochemistry, photochemical charge separation, exciton coupling, Circular Dichroism, Magnetic Circular Dichroism, fluorescence, electrochromic shift

## Abstract

Far-red light (FRL) Photosystem II (PSII) isolated from *Chroococcidiopsis thermalis* is studied using parallel analyses of low-temperature absorption, circular dichroism (CD) and magnetic circular dichroism (MCD) spectroscopies in conjunction with fluorescence measurements. This extends earlier studies (Nurnberg *et al* 2018 Science 360 (2018) 1210-1213). We confirm that the chlorophyll absorbing at 726 nm is the primary electron donor. At 1.8 K efficient photochemistry occurs when exciting at 726 nm and shorter wavelengths; but not at wavelengths longer than 726 nm. The 726 nm absorption peak exhibits a 21 ± 4 cm^−1^ electrochromic shift due to formation of the semiquinone anion, Q_A_^•-^. Modelling indicates that no other FRL pigment is located among the 6 central reaction center chlorins: P_D1_, P_D2_ Chl_D1_, Chl_D2_, Pheo_D1_ and Pheo_D2_. Two of these chlorins, Chl_D1_ and P_D2_, are located at a distance and orientation relative to Q_A_^•-^ so as to account for the observed electrochromic shift. Previously, Chl_D1_ was taken as the most likely candidate for the primary donor based on spectroscopy, sequence analysis and mechanistic arguments. Here, a more detailed comparison of the spectroscopic data with exciton modelling of the electrochromic pattern indicates that P_D2_ is at least as likely as Chl_D1_ to be responsible for the 726 nm absorption. The correspondence in sign and magnitude of the CD observed at 726 nm with that predicted from modelling favors P_D2_ as the primary donor. The pros and cons of P_D2_ vs Chl_D1_ as the location of the FRL-primary donor are discussed.

**TOC Graphic:** 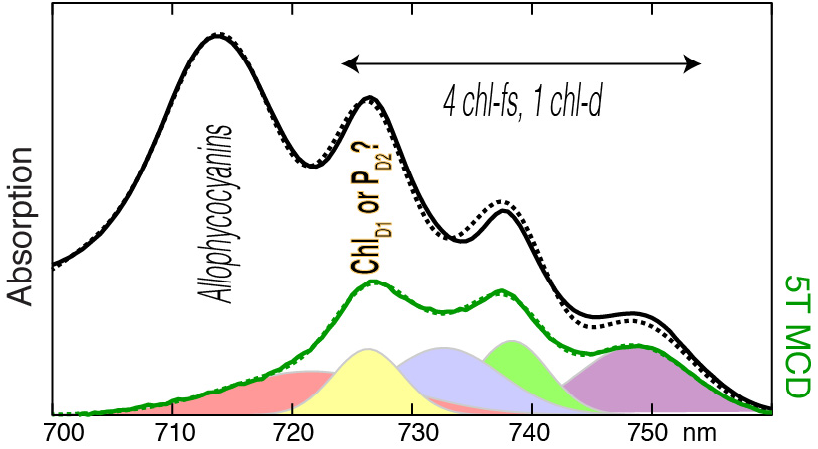

**Highlights:**

- Primary Donor confirmed at 726 nm
- Determination of far-red chl pigment Q_y_ excitation positions, widths, CD and MCD amplitudes
- Quantification of electrochromic shifts and Q_A_^•-^ photoconversion yield
- Electrochromic shift consistent with primary donor at either Chl_D1_ or P_D2_
- The CD amplitude favors the primary donor at P_D2_

## 1 INTRODUCTION

Oxygenic photosynthesis in the cyanobacteria *Chroococcidiopis thermalis*, when grown in far-red light (FRL), is based on photochemistry occurring on *chl-f* or *chl-d* [1]. This constitutes an unexpected new paradigm in photosynthesis. Photosystem II core complexes (FRL-PS II) of *C. thermalis* contain 4 *chl-f* molecules and 1 *chl-d*, whilst the majority (~30) of the chlorin pigments remain as *chl-a*.

The original spectroscopic studies of the isolated FRL-PS II provided a model in which one of the long wavelength pigments (FRL-chl) played the role of the primary donor, while the 4 remaining FRL-chls served as long wavelength antennas [1]. The primary donor was identified [1] as Chl_D1_ based on the following evidence and arguments: a) trapping of Pheo ^•-^ at room temperature resulted in electrochromic changes at 720 nm and the apparent loss of the usual dominant blue-shift at ~ 685 nm attributed to *chl-a* Chl_D1_; b) Q_A_^•-^ formation at low temperature resulted in electrochromic changes at 725 nm and again the apparent loss of the usual dominant blue-shift at ~ 685 nm attributed to the *chl-a* Chl_D1_; c) The presence of amino acid sequence changes in the FRL-D1 protein conserved in all known FRL-D1 sequences, which were modelled as potential H-bond donors to the ring I formyl group of either *chl-d* or *chl-f* in the Chl_D1_ location [1]; d) in *Acaryochloris marina,* a similar bandshift was seen (as well as similar H-bonding amino acids) already attributed to *chl-d* at the Chl_D1_ location [2]; e) functional arguments based on Chl_D1_ taking the primary donor role in canonical *chl-a*-PSII (see e.g. [3–7]); f) The chemical properties of *chl-f* [8] and the redox properties [9] of both *chl-d* and *chl-f* favor its location on Chl_D1_ (see below and [1]).

The reaction center of c*hl-a-*PS II contains 2 *pheo-a* and 6 *chl-a* molecules. P_D1_ and P_D2_ have partial ring overlap, forming a chlorophyll pair that is the counterpart of the “special pair” in purple bacterial reaction centers. The P_D1_/P_D2_ exciton coupling term in PS II [10] at 158 cm^−1^ is far smaller than the ‘special pair’ coupling of ~900 cm^−1^ determined for *Rhoropseudomonas viridus [11]*. Chl_D1_ and Chl_D2_ are counterparts of the monomeric bacteriochlorophylls, B_L_ and B_M_, of the bacterial reaction center; they are sometimes anomalously referred to as “accessory pigments” despite the demonstrated role Chl_D1_ (and B_L_) has in primary charge separation [3]. The two *pheo-a* pigments, Pheo_D1_ and Pheo_D2_, are also core pigments, with Pheo_D1_ having a key electron acceptor role. ChlZ_D1_ and ChlZ_D2_ are not part of the central group of 6 chlorins as they are peripheral and mainly play light-harvesting roles, although ChlZ_D2_ is part of a side-path electron donation system that works at low quantum yield under some circumstances [12, 13]

P_D2_ was not considered as a candidate for the primary donor role in FRL-PS II although it is oriented and located in a way that can give rise to an electrochromic shift very similar to that calculated for Chl_D1_. Here we examine the possibility that P_D2_ is the primary electron donor in FRL-PS II.

## 2 Materials and Methods

FRL-PS II samples were prepared as described in [1].

All spectra were taken on a single, purpose-built multifunctional apparatus [14] built around a 0.75 M Czerny-Turner spectrometer (Spex) and a superconducting magnet cryostat (Oxford Instruments SM-4). The system is able to simultaneously record absorption, CD and MCD spectra at temperatures between 1.8 K and 300 K. This system also can be rapidly reconfigured to measure emission and excitation spectra, without disturbing the cryogenic sample.

The apparatus has a sample handling system [15, 16] that minimizes exposure of the sample to oxygen and ambient light, whilst allowing rapid cooling of a room temperature sample to cryogenic temperatures under controlled conditions. Samples were visualized with a near-IR light (>780 nm) sensitive camera at either room temperature or at cryogenic temperatures. The presence of bubbles, cracks or defects in the prepared sample could be immediately determined without inducing any photoconversion of a dark-adapted sample.

The PSII samples contained cryoprotectant glycerol/ethylene glycol (1:1) at a final concentration of 45 % (v/v). Prior to dark adaption, samples were loaded under low light into specially designed [16] strain-free quartz optical cells of 500 μm, 200 μm or 50 μm pathlength. The PSII samples were then dark-adapted for 5 min before being rapidly cooled.

The light source for the absorption measurements was a highly stabilized 250 W quartz-halogen lamp. The fluence of the measurement light at the sample was reduced to a point that absorption measurements did not induce a significant degree of Q_A_ reduction in the PS II sample. This was established by determining that there was no significant difference between two successive absorption measurements. The necessary reduction in measurement light seen by each PS II assembly involved defocusing the measurement light over a large area (~8 mm diameter) sample and narrowing the slit widths of the monochromator, whilst using a large area high quantum efficiency Hamamatsu R 669 photomultiplier to detect the transmitted light [17]. Electrochromism was induced with a highly stabilized 250 W quartz-halogen lamp passed through either a green bandpass filter combined with a 10 cm pathlength water filter to provide 1-2 mW of broad spectrum green light, or through an optical system [14] based on a Spex Minimate 0.25 M double monochromator, providing 6-8 μW of light in the 600 nm - 760 nm range and having a bandwidth of 2.4 nm FWHM (see Fig. S2), or a Schwartz EO Ti:S laser pumped with a 532 nm Continuum Verdi G.

CD and MCD measurements required much higher measurement light levels. It is not possible to measure useful CD or MCD spectra and maintain the sample in the Q_A_ state [18]. Thus, samples were illuminated with green light (300 s at 1-2 mW/cm^2^) forming the Q_A_^•-^ state *prior* to the measurement [19] to minimize actinic effects during the course of CD and MCD measurements.

Emission was excited with the highly stabilized lamp and Spex Minimate system described above and then detected through the same 0.75 M Czerny-Turner spectrometer as the absorption with a Hamamatsu R 943-02 photomultiplier tube.

Data were analyzed using IGOR pro software and calculations were made using MATLAB.

## 3 Results and Discussion

### 3.1 Canonical chl-a-PS II and FRL-PS II comparisons

Fig. 1 provides a comparison of low temperature spectra of FRL-PS II core complexes of *C. thermalis* with those of *T. vulcanus*. The latter is a well-determined *chl-a* containing PSII, i.e. *chl-a-*PS II. FRL-PS II exhibits a number of well-resolved absorption features in the 700-760 nm range that are not present in *chl-a*-PS II, as previously reported [1]. Fluorescence spectra of the two systems are also significantly different (Fig. 1 and SI, Fig. S3 of [1]), with *chl-a*-PS II fluorescence peaking near 690 nm and the FR-PS II peaking near 752 nm.

**Fig. 1.**
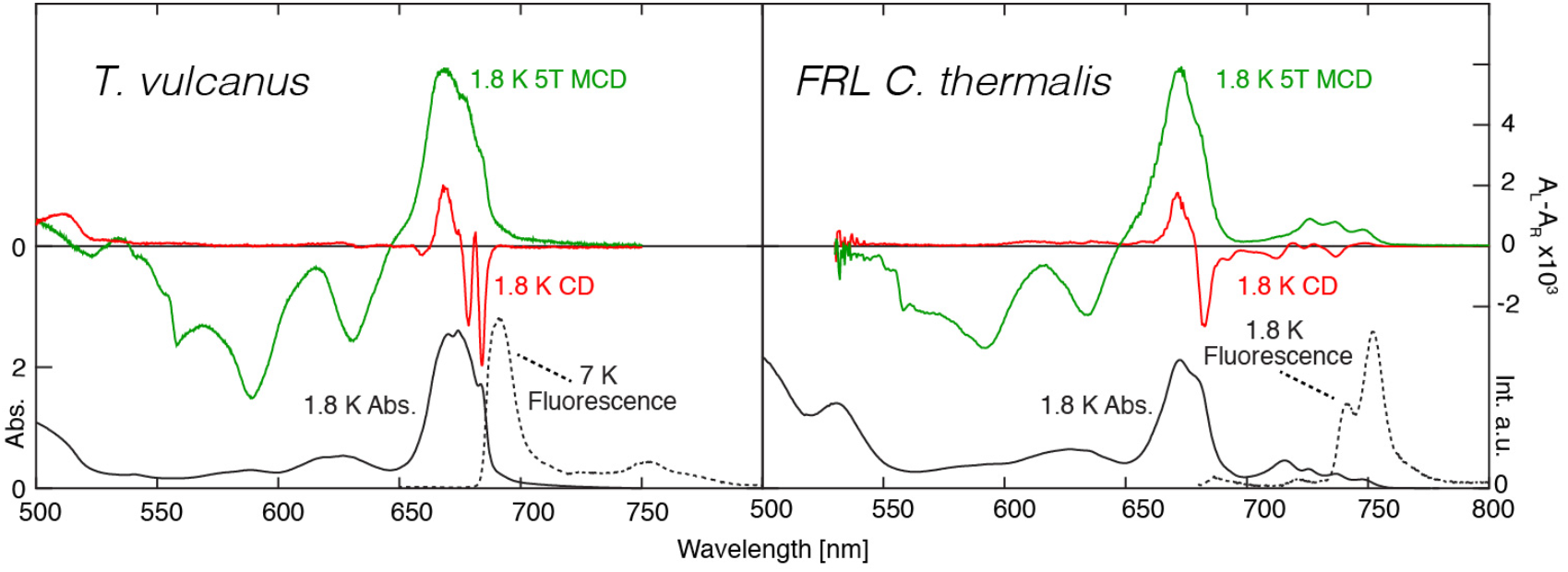
Low temperature absorption, CD, MCD and fluorescence spectra (as indicated) of chl-a-PS II from T. vulcanus (left, adapted from reference [20]) and FRL-PS II of C. thermalis (right). Corresponding spectra are set to the same integrated Q_y_ absorption strengths.

Absorption features in the region beyond ~720 nm involve Q_y_ excitations of 4 *chl-f* molecules and one *chl-d.* Absorption integrals (see SI-1, Fig. S1) are consistent with there being ~30 *chl-a* molecules, 2 *pheo-a* pigments and 5 far-red chlorophylls in FRL-PSII, in agreement with biochemical analyses [1].

### 3.2 Identification of the primary donor

Fig. 2 shows that 735 nm illumination at 1.8 K induces a *minimal* electrochromic shift amplitude near 726 nm. At 77 K, illumination at 750 nm leads to a minimal shift amplitude whereas illumination at 735 nm leads to a small but significant shift amplitude. The small shift at higher temperatures is attributable to the optically excited lower energy pigment (peaking at 738 nm) leading to charge separation by way of thermally activated access to the primary donor at 726 nm. The energy gap between peaks at 726 nm and 738 nm is 12 nm (220 cm^−1^). This corresponds to ~4 kT at 77 K pointing to a possible thermally activated primary donor occupation of ~2 %. We are thus able to confirm that the 726 nm peak is that of the primary donor [1].

**Fig. 2.**
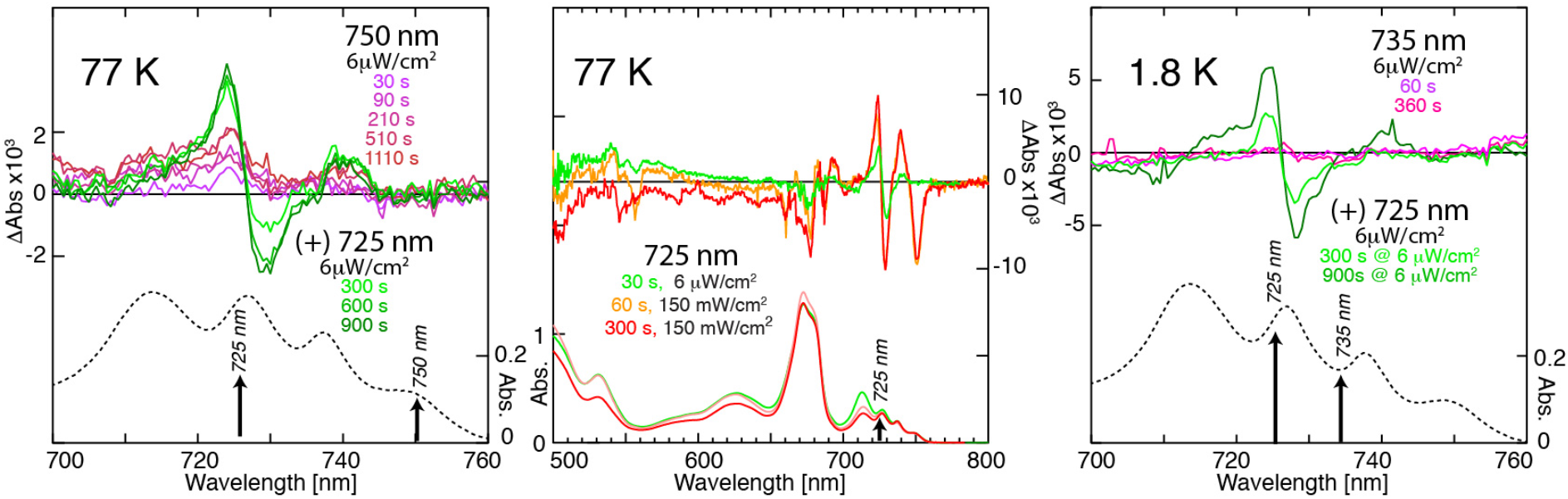
The left panel shows shift and absorption spectra measured at 77 K, for low power illuminations at 750 nm followed by additional low power illuminations at 725 nm. The central panel compares the 77 K spectrum for a sample illuminated for 30 s with low power 6 μW/cm^2^ 725 nm illumination (green), followed by illumination by a 1500 times more powerful, 150 mW/cm^2^ Ti:S laser source, also at 725 nm for 60 s and 300 s (orange and red traces respectively). The right panel shows difference features induced by illuminating at 1.8 K with low power 735 nm radiation, followed by low power 725 nm illumination.

At high excitation fluences, achieved by using a Ti:S laser at 725 nm, additional features are generated in difference spectra (central panel in Fig. 2). Most apparent is a bleach at 749 nm, along with an absorption increase in the 738 nm region. The bleach at 749 nm is associated with partial oxidation of the lowest energy FRL-chl. Such photo-oxidation processes are known [17, 21]. The peak position of the increased intensity component is at 739.5 nm and does not match the peak position of the band seen in absorption. A blue-shift of this type is characteristic of non-resonant photophysical hole-burning [22–24] although such processes are more pronounced at lower temperatures.

### 3.3 Variation in allophycocyanin content of FRL-PS II and Global Analysis

Absorption at wavelengths longer than 700 nm is attributed primarily to FRL-chl. However, there is significant absorption at the shorter wavelength end of the 700 nm to 740 nm region that can be attributed to allophycocyanins [1]. More highly purified FRL-PS II preparations show that the absorption feature near 715 nm becomes depleted, whereas the three other distinct features are retained. Fig. 3 shows FRL-PS II spectra of samples with different allophycocyanin content.

**Fig. 3.**
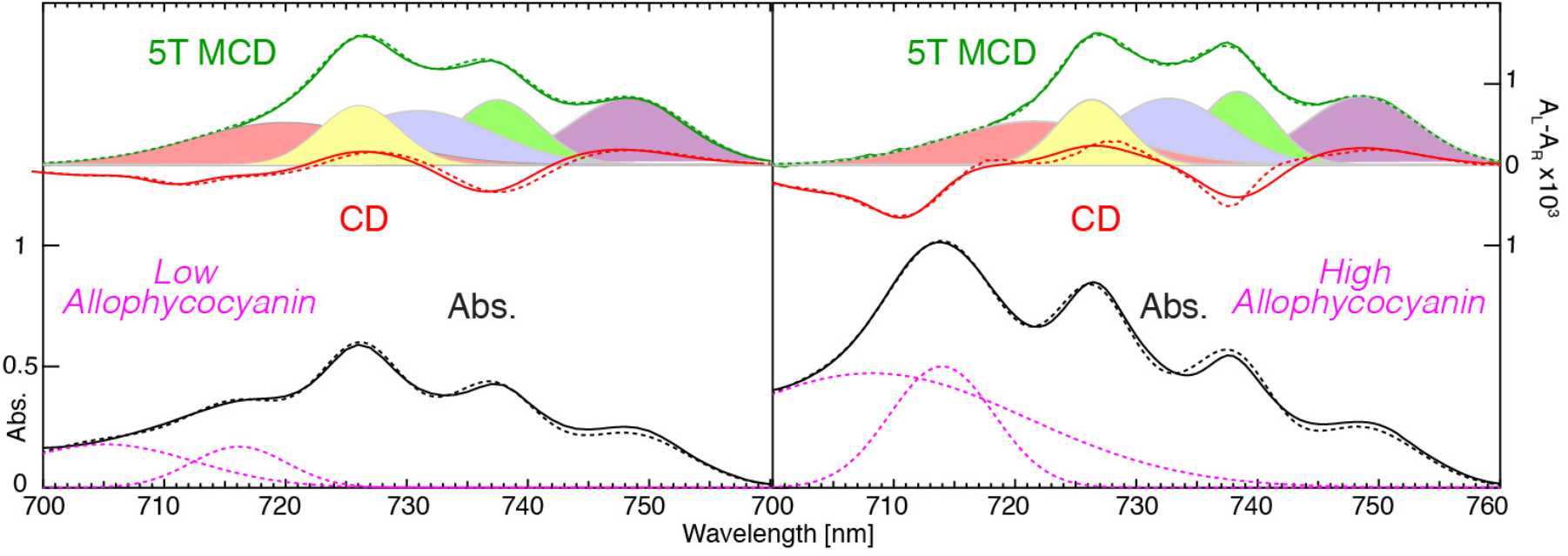
Absorption (solid black traces), MCD (solid green traces) and CD (solid red traces) of high and low allophycocyanin containing FRL-PS II preparations, scaled to the same area in MCD. A systematic analysis of the MCD (see text and SI-3) leads to the quantification of the 5 individual FRL-chl Q_y_ MCD components shown in Table S1. The MCD contribution of these 5 Q_y_ excitons are shown under the MCD spectrum in the figure as coloured overlapping Gaussians. The primary donor (at 726 nm) is yellow. The subsequent fit of absorption spectra (dashed black traces) and CD spectra (dashed red traces), using the positions and widths of the 5 components as determined from the MCD analysis (see text and Table S1) closely follows the experimental data. Allophycocyanin absorption components are estimated (see SI-1) as the dashed magenta curves.

The 700 nm to 715 nm region of FRL-PS II shows little MCD, but significant CD and absorption. The absorption and CD intensity in this region vary strongly with allophycocyanin content (Fig. 3), showing it to be dominated by allophycocyanins. The fluorescence spectra of high-allophycocyanin FRL-PS II (Fig. 1 and SI-2, Fig. S2) exhibit a weak peak near 730 nm, associated with emission from allophycocyanins, indicating that they are, at least partially, disconnected from PSII.

There are three well-resolved peaks in FRL-PSII spectra at wavelengths longer than 720 nm that are not due to allophycocyanins. These are due to the FRL-chls. An analysis of high and low allophycocyanin FRL-PSII samples (Fig. 3, Table S1 and SI-3) establish small changes in the position and width of these peaks, indicating only a minor disturbance of the FRL-PS II proteins associated with the purification process.

The spectral position and width of all 5 FRL-chl Q_y_ excitations were determined by a parallel, information-driven, global analysis of simultaneously measured absorption, CD and MCD spectra, such as those seen in Fig. 3. The MCD spectra are key to this analysis as they have no significant contribution from overlapping signals from allophycocyanins. Another important characteristic of the FRL-chl Q_y_ excitations is the MCD amplitude relative to absorption ratio B/D which in turn is proportional to ΔA/A at a specific applied magnetic field. This ratio is known to vary and also to be indicative of the role that the pigment takes [18, 25, 26]. The primary donor, in particular, exhibits a narrow Q_y_ exciton for which its MCD B/D value is significantly reduced (by ~50%) compared to pigments involved in light harvesting, which are less strongly coupled to other chlorophylls [18, 27]. All purely electronic excitations (which include exciton coupled Q_y_ bands) that exhibit B term-type MCD have line-shapes that follow the absorption profile of each exciton component [28–30]. Each FRL-chl is taken to have the same integrated Q_y_ absorption intensity. Exciton coupling-induced intensity re-distribution (Section 3.3) is thus considered to be minor. The oscillator strength differences between *chl-a*, *chl-d* and *chl-f* have been shown [31] to be relatively small. In contrast, the CD amplitude of each Q_y_ exciton component band can vary in both magnitude and sign, and is determined by exciton coupling to neighboring pigments [32]. For the purpose of this analysis, the CD bandshapes were assumed to be the same as that of the absorption and MCD. However, CD line shapes may follow those of their absorption features less closely when vibrational sideline features become involved [26, 33]. The overall fit to the CD shown in Fig. 3 is less precise than that of the absorption and MCD. This may arise partly as a result of the influence of vibrational sideline CD. Attempts to fit the CD spectrum in the 726 nm region with the CD component of the primary donor constrained to a small relative amplitude were unsuccessful. The positive CD seen in this region is associated with the primary donor (see SI-3 and Fig. S4).

A description of the global analysis is provided in SI-3. Table S1 provides the relevant output. Very similar positions and widths for each FRL-chl Q_y_ MCD band were obtained from both the low and high allophycocyanin samples shown in Fig. 3. There is a variation of relatively strong CD features associated with allophycocyanins present in the high and low allophycocyanin FRL-PS II samples which contributes to uncertainties in determining the CD of the broader FRL-chl Q_y_ excitons. However, both the sign and magnitude of CD for each of the three narrow FRL-chl Q_y_ peaks remains well determined. This is considered in detail in SI-3. A graphical presentation of the individual CD components, as seen for the MCD components in Fig. 3 is given in Fig S4.

### 3.4 Analysis of *Electrochromic Shifts*

Illumination of PS II core complexes at cryogenic temperatures induces metastable spectral shifts. Prominent shifts are attributed to the formation of Q_A_^•-^. This radical anion is adjacent to Pheo_D1_ and close to Chl_D1_. It induces relatively large electrochromic shifts of the Q_x_ band of Pheo_D1_ (historically named the C550 shift) [34] and of the Q_y_ band of Chl_D1_ [23, 35, 36]. Fig. 4 shows both absorption and illumination difference spectra for a FRL-PS II sample.

**Fig. 4.**
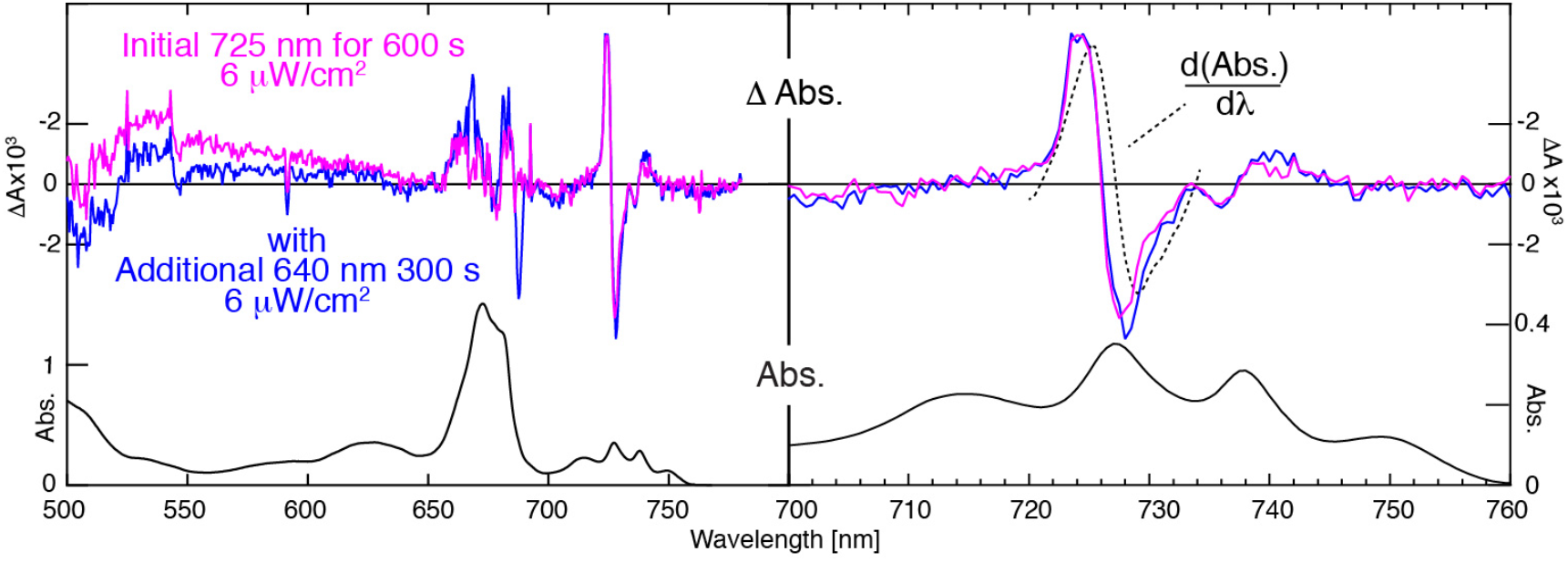
Lower left panel: 1.8 K absorption spectrum (black line). Upper left panel: Difference spectra after 1. illumination at 725 nm for 600 s with an incident fluence of 6 μW/cm (magenta line), and 2. the total change in absorption induced by the first illumination plus additional illumination at 640 nm (blue line). Right panel: Long-wavelength detail of the absorption and difference spectra shown in the left-hand panels compared to the (scaled) differential of the absorption spectrum taken before illumination (dashed line). This data is analyzed in more detail in SI-4 and Fig. S6.

These spectra are interpretable in terms of previous analyses of PS II core complex electrochromism [23, 36]. A comparison of representative electrochromism spectra in PS II core complexes of 4 organisms is shown in Fig. S5. In FRL-PS II the C550 shift of the Pheo_D1_ Q_x_ feature at 544.5 nm is close to the position in other cyanobacterial PS II systems [23]. In some FRL-PSII preparations (SI, Fig S5) a bleach upon photoreduction of Q_A_ identifies (the initially reduced) cytb559 absorption in FRL-PSII to be at 556.5 nm. The dominant blue-shift of Q_y_, seen at 1.8 K near 683 nm for *chl-a*-PS II [17, 37] now appears near 726 nm under similar conditions, as reported earlier [1]. This is consistent with the 726 nm band being associated with the primary donor. In *chl-a*-PS II, the Q_y_ electrochromic shift pattern is dominated by the Pheo_D1_ and Chl_D1_ excitons. The site-energies of the Chl_D1_ and Pheo_D1_ pigments are similar and there is a significant exciton matrix element between them (Table S3). When pigment excitations are close in energy, this matrix element leads them to interact strongly, with intensity being transferred between the resulting excitations [23]. Calculation of the electrochromism requires independent exciton computations of both Q_A_ and Q_A_^•-^ configurations of PS II. Electrochromic effects displace site energies significantly, causing the mixing of exciton states to change. The situation with FRL-PS II is different, as the primary donor is displaced to lower energy by ~ 800 cm^−1^ or more from the *chl-a* excitations. Thus, the effects of the coupling matrix elements between the primary donor and spatially adjacent pigments such as Pheo_D1_ and P_D1_ are reduced by the energy differences between them, although the matrix elements themselves do not change.

The spectral isolation of the primary donor Q_y_ absorption in FRL-PS II assists in the estimation of both its electrochromic shift and the yield of Q_A_^•-^ formation. The observed shift appears as a significant fraction of the linewidth and the absorption difference spectrum does not cross precisely at the maximum of absorption (Fig. 4) as would be the case for electrochromic shifts much smaller than the linewidth. The analysis is presented in SI-4 and leads to a yield of 5.5 ± 0.5 % and an initial estimate of the shift at 42 ± 4 cm^−1^.

However, a quantitative comparison of electrochromic shift data accumulated at 77 K and 1.8 K data taken at higher fluences with low fluence 1.8 K data (Fig. 2), point to a shift magnitude of 42 cm^−1^ to be an overestimate. Instead, it seems that the pigments that photoconvert with high efficiency are only a subset of the total pigment population absorbing at 726 nm. The subset is narrower than the total distribution and also shifted (by ~1 nm) from the peak maximum as measured in absorption. The complete analysis is presented in SI-4 and provides a value for the electrochromic shift of 21 ± 4 cm^−1^.

There is no observable electrochromic shift of the lowest energy feature at 749 nm. The narrow negative feature peaking at 687.5 nm seen in Fig. 4 is not induced by illumination at 725 nm but only with higher energy (640 nm) illumination. This feature is attributed to electrochromism of *chl-a*-PS II present in the FRL-PS II sample as a contaminant. The data in Fig. 2 and Fig. 4, along with the discussion in SI-4 establish that low power 725 nm illumination does not induce significant *chl-a*-PS II photochemistry, i.e. does not induce Q_A_^•-^ formation as measured by the characteristic Chl_D1_ electrochromic shift in the 685 nm region.

There is a smaller red shift of ~7 ± 2 cm^−1^ for the 738 nm peak. This feature is not present at the lowest illumination fluences, see Fig. 2. and Fig. S6. There are also shoulders near the main shift feature that become more pronounced at higher fluences. These may be attributed to secondary photophysical and photochemical processes of the type that also occur in *chl-a*-PS II and related systems (see SI-4).

The yield of Q_A_ ^•-^ compared to that seen for most other organisms remains low even at high fluences (see SI-4, Fig. S5). However, yields are also very low in *Acaryochloris marina* [38], a system in which PS II is dominated by the FRL-chl *chl-d*. Strong illumination of FRL-PS II (SI-4, Fig. S6) leads to increased yields, but also induces additional photochemical and photophysical changes [17].

### 3.5 Exciton Calculations

The first widely accepted multi-spectral model [39] for the reaction center of *chl-a*-PS II used a range of spectral data as input. Exciton couplings between pigments were based on the crystallographically determined distances between chlorins and also the relative orientation of their transition dipoles. A range of exciton realizations were generated to account for the non-correlated inhomogeneity of pigment site-energies. Inhomogeneous widths were seen to be comparable to exciton coupling energies. The authors used a Redfield model for coupled Brownian oscillators to account for temperature dependent data. A simplified version of this model [23], restricted to the analysis of low temperature data, was used to calculate the absorption, CD and electrochromic shifts of the intact reaction center within PS II core complexes in a range of organisms. In this work [23], calculations of electrochromic shifts as well as CD changes associated with both Q_A_^•-^ formation and Pheo_D1_ reduction were made.

In the current work, the simplified model [23] is adapted to model the FRL-PS II reaction center absorption spectra, CD and electrochromic shifts. Protein structure, coupling matrix elements, transition dipoles and static dipoles were taken to be the same as for *chl-a*-PS II.

Firstly, the previously reported analysis [23] of the Q_A_^•-^-induced Q_y_ electrochromic shift pattern, as calculated for plant PS II core complexes at 1.7 K [17], was reproduced and refined (left panel of Fig 5). Then, using the same model, the essential features of the experimental electrochromic pattern in FRL-PS II were reproduced. This model does not take into account i) electrochromic shifts from the low quantum yield side-pathway of electron transfer, including ChlZ bleaching near 670 nm [36], and ii) photophysical shifts of the lowest energy states of CP47 [24] and CP43 [33] near 690 nm. The lowest excited states [40] of peripheral antennae are long-lived and prone to non-resonant hole-burning upon sustained illumination. This last process leads to a redistribution of inhomogeneously distributed pigment site energies to higher energies and in particular, to a reduction of absorption intensity in the lowest energy 690nm to 700 nm region of spinach PS II, as seen in Fig. 5.

**Fig 5.**
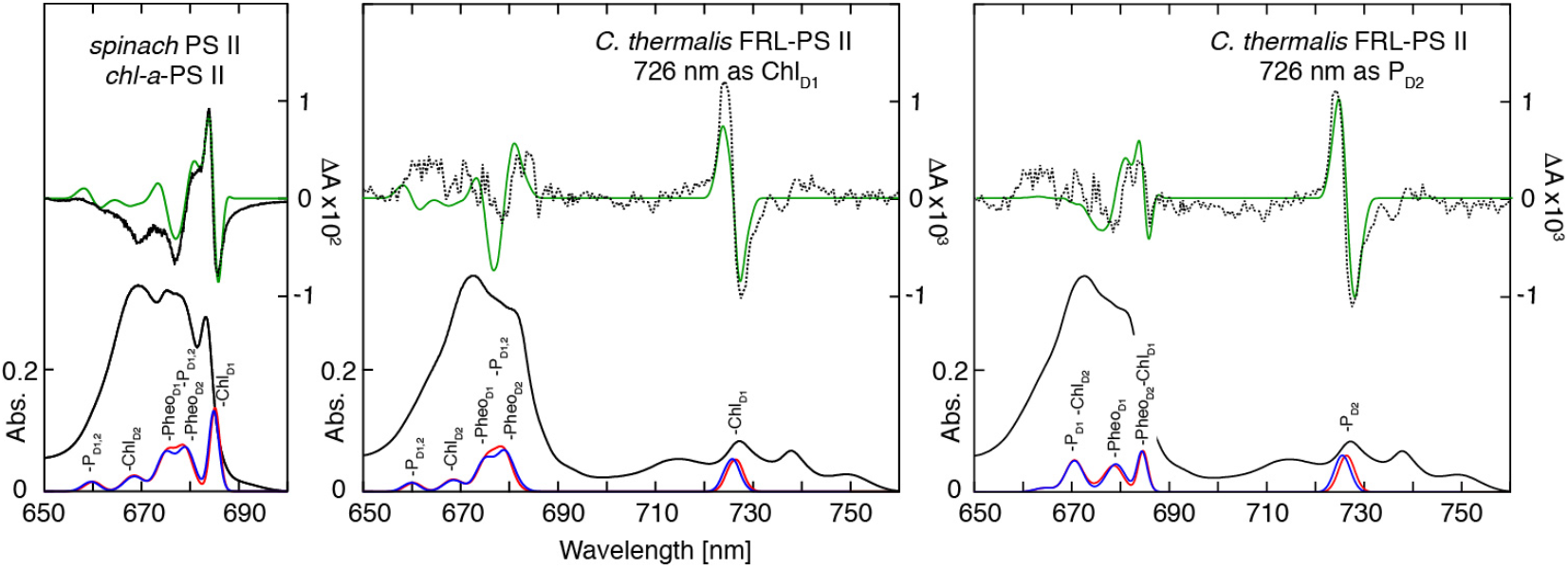
Lower black traces are absorption spectra of spinach chl-a-PS II core complexes [17] (left panel) and FRL-PS II (center and right panels) at 1.8 K. The upper black traces are experimental electrochromic difference spectra, for 85 % conversion of Q_A_ to Q_A_^•-^ for spinach and ~8% conversion for FRL-PS II (See SI-3). The corresponding green curves are the modelled electrochromism spectra for spinach core complexes (left panel) and FRL-PS II using the Chl_D1_ or P_D2_ options for the location of the 726 nm pigment. The lower red and blue curves are the corresponding calculated absorption spectra of the 6 inner reaction center chlorins for the Q_A_ and Q_A_^•-^configurations respectively. The upper green traces show the calculated electrochromism. Site energies and the exciton coupling matrix used are given in the Tables S2 and S3.

The amplitude of the Q_A_^•-^ induced electrochromism reported in spinach PS II core complexes [17], as reproduced in Fig. 5, was that for a Q_A_^•-^ yield measured as ~85%. The knowledge of both the yield and amplitude allows an independent and experimental determination of the value of Δμ/f, where Δμ is the change in static electric dipole moment between the ground and excited state and f is the dielectric correction factor. Using the previous [41, 42] value of Δμ= 1.8 D for *chl-a* and 1.35 D for *pheo-a*, and taking f = 3, as used earlier [23], an excellent fit to the reported [17] spinach PS II shift amplitude near 685 nm is obtained, as shown in Fig. 5. This confirms the use of a value of f=3.

The values of Δμ=1.8 D for *chl-a* and 1.35 D for *pheo-a* with f = 3 used to fit spinach PS II were used to model FRL-PS II. Except for the FRL 726 nm pigment, for which the position and linewidth was taken from Table S1, site energies and widths of the other (*chl-a*) pigments were left unchanged. The calculations shown in Fig. 5 show that with a (single) FRL-chl placed in the reaction center, either at the Chl_D1_ or P_D2_ position, a strongly blue shifted FRL-PS II electrochromic feature appears (see also Fig. S9). Placing more than one FRL-chl in the reaction center leads to a number of prominent shift features in the far-red region of the electrochromic pattern. Only one prominent shift feature is seen in the low fluence spectra (Fig. 4, Fig. 5). This provides evidence that there is only one FRL-chl in the reaction center. This more detailed treatment confirms the interpretation in the original study [1].

When comparing the calculated electrochromic shift spectra with Chl_D1_ or alternatively P_D2_ as the primary donor against experimental data (Fig. 5), both options reproduce the sign and amplitude of the shift seen at 726 nm. The P_D2_ option provides a somewhat better overall fit to the electrochromic pattern in the *chl-a* Q_y_ region from 650 nm to 690 nm. FRL PS II data in this region suffers from low signal-to-noise due to i) the higher absorbance present in the *chl-a* Q_y_ spectral region, ii) the requirement of minimal actinic light and iii) the relatively low yield of Q_A_^•-^ formation in FRL-PS II (see Fig. S6) compared to spinach *chl-a*-PS II.

The calculated amplitude of the 726 nm Q_A_^•-^ shift is slightly smaller for the Chl_D1_ primary donor case than is for the P_D2_ case, despite P_D2_ being further away (2.6 nm vs 2.2 nm) from Q_A_. This is a consequence of the orientation of the electric field with respect to Δμ leading to a larger site-energy shift for P_D2_ (Table S4). The chlorin Q_y_ site energy shifts for Q_A_^•-^ formation (Table S4) shows that the P_D1_ and P_D2_ shifts (5.8 cm^−1^ and 18 cm^−1^) are comparable to the Chl_D1_ shift (14.4 cm^−1^).

Exciton modelling can be used to simultaneously calculate the lineshape and amplitude of both CD and absorption spectra [26, 32] without the introduction of any additional parameters. The amplitude of the CD calculated in this way provides a useful internal check of exciton calculations. For the isolated reaction center preparation RC-6 both absorption and CD line-shapes have been successfully modelled [39]. A quantitative comparison of CD and absorption amplitudes was not made at that time. A detailed comparison of more recent, simultaneously taken, lower-temperature absorption and CD spectra, of both RC-6 and RC-5 [26] show that the calculated CD line-shape as well as amplitude are in agreement with the exciton calculations. The calculations done for RC-6 and RC-5 use the same exciton matrix elements (Table S3) as those used here. The only difference in the former calculations were that the linewidths and site-energies introduced were adapted to take into account the increased line-broadening and site-energy blue shifts present in isolated reaction centers such as RC-6, compared to PS II core complexes.

Isolated FRL-PSII reaction centers have not yet been prepared. Consequently, it is only possible to compare theoretical and experimental CD data for the single FRL-chl present in the reaction center component of the FRL-PS II core complex assembly, i.e. the pigment absorbing at 726 nm. The CD characteristics of the other *chl-a* and *pheo-a* FRL-PS II reaction center pigments, i.e. those absorbing in the 650 nm to 690 nm region, are obscured by the dominating CD of *chl-a* present in CP43 and CP47 peripheral antennas (Fig. 1). A more complete analysis would look to calculate the exciton CD of the entire FRL-PS II core assembly but this is not at present feasible. Exciton CD calculations that are restricted to the reaction center pigments of FRL-PS II are a reasonable starting point as couplings between reaction center pigments and pigments in the CP43 and CP47 peripheral antennae are inherently weaker than the (relatively strong) couplings between reaction centre pigments (Table S3). This is due to the greater inter-pigment distances involved.

Fig. 6 demonstrates that the computed absorption and CD spectra of the 6 reaction center chlorins in FRL-PS II, shown as coloured green and red profiles respectively, vary significantly depending on whether the primary donor is located at Chl_D1_ or P_D2_. Of the 6 reaction center chlorins, only the FRL-chl primary donor is directly accessible experimentally, because it has CD and absorption features that are not buried under the absorption peak arising from ~30 *chl-a* and *2 pheo*-a pigments in the 650 nm - 690 nm region.

**Fig 6.**
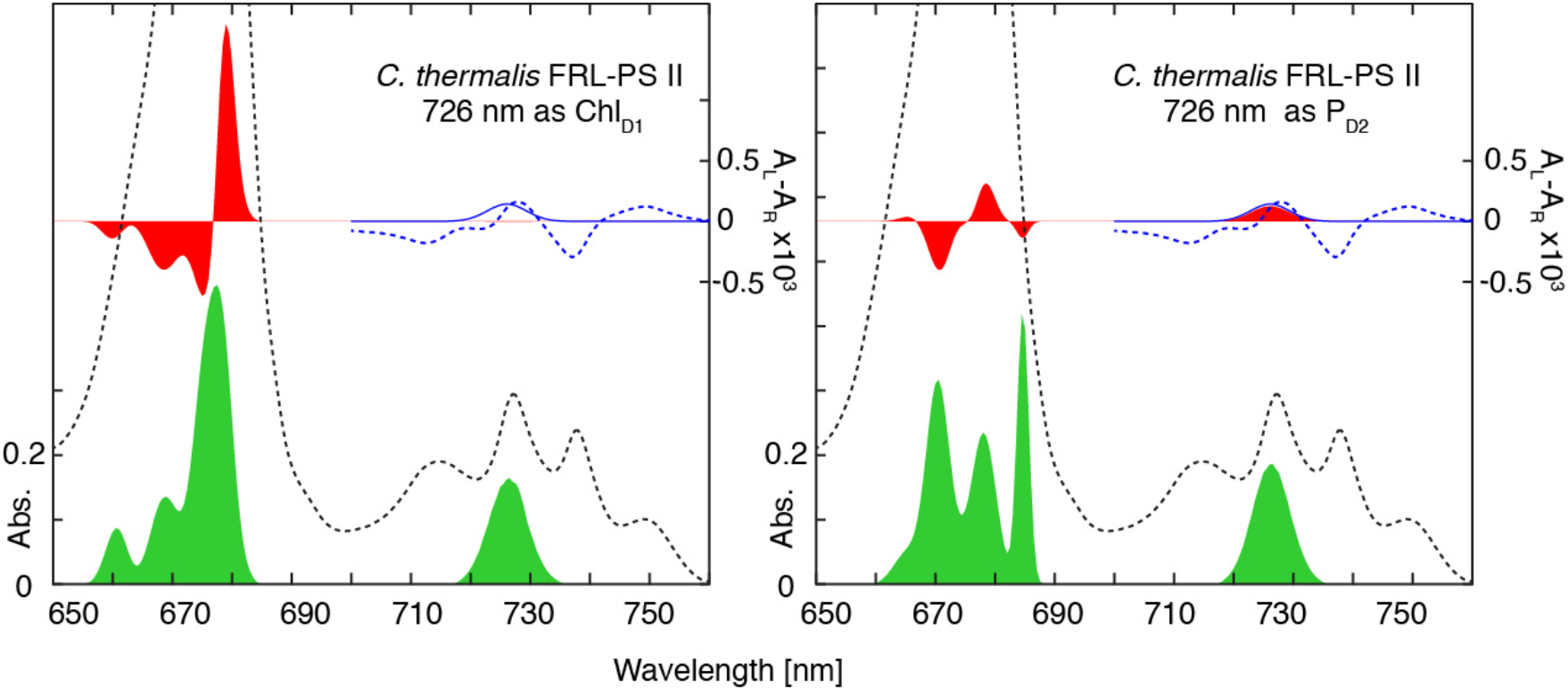
For both panels, the dashed black trace (left-hand scale) is the 1.8 K absorption spectrum of a FRL-PS II preparation. The dashed blue trace is the simultaneously-recorded CD spectrum in the red region (right-hand scale). The solid green profiles are the exciton-modelled absorption spectra (see text) of the inner reaction center chlorins of FRL-PS II and the red profiles are the corresponding calculated CD spectra, with either Chl_D1_ or P_D2_ as the primary donor as indicated. The solid blue trace shows the CD component of the primary donor at 726 nm. The position, width and amplitude of the primary donor CD at 726 nm (Table S1) was determined by the global fitting procedure described in the text. Experimental FRL-PS II absorption and CD spectra are scaled to the calculated (reaction center only) spectra by assuming that the integrated absorption area over the entire Q_y_ region corresponds to a total of ~35 chl and 2 pheo-a pigments with the reaction center having 6 chl and 2 pheo-a pigments. Integrals are shown in Fig S1.

The sign and amplitude of the calculated primary donor CD agrees well with the (positive) experimental CD value when P_D2_ is the primary donor. When Chl_D1_ is the primary donor, the calculated CD is barely visible. It is 30x smaller and is calculated to have a negative sign. The CD spectrum of a multi-chromophore array such as a PS II reaction center arises from a complex interplay between exciton coupling matrix elements (Table S3), site energies (Table S2) and relative orientations of transition dipoles of the interacting chromophores. Thus, it may not be immediately interpretable. However, there is only one FRL-chl in the reaction center of FRL-PS II (i.e. the pigment at 726 nm) and the Q_y_ of this pigment is displaced by ~1000 cm^−1^ from the other pigments (*chl-a* and *pheo-a*) in the reaction center. This displacement is larger than the values of the exciton coupling matrix elements shown in Table S3. With coupling matrix elements and orientations being held constant, a large red-shift of the chromophore will tend to have the effect of reducing its CD, whether it be located at Chl_D1_ or P_D2_.

A quantitative measure of the strength of CD is the parameter ρ_A_, the ratio of CD amplitude to absorption, ΔA/A. The CD of the *Chl-a* exciton region at 680 nm in RC-6 is experimental and theoretically evaluated as ρ_A_ = 1.2 × 10^−3^. This ρ_A_ value is significantly less than the calculated CD of the Chl_D1_ exciton itself which has ρ_A_ = 2.5 × 10^−3^ [26]. The reduction in ρ_A_ is due to the significant overlap of the Chl_D1_ exciton with nearby excitons having the opposite CD sign. In FRL-PS II the primary donor exciton is well separated from other reaction center excitons and such cancelling does not occur. Thus, we can directly compare the experimental ρ_A_ of the primary donor in FRL-PS II with the calculated value of ρ_A_ = 2.5 × 10^−3^ for the Chl_D1_ exciton in *chl-a*-PS II.

From our global analysis of FRL-PS II data, the CD of the primary donor in FRL-PS II at 726 nm has ρ_A_ = 0.5 x10^−3^. The calculated values for the primary donor CD ΔA/A at 726 nm is ρ_A_ = 0.6 × 10^−3^ when P_D2_ is the primary donor and −0.02 × 10^−3^ when Chl_D1_ is the primary donor. In both cases, the calculated ρ_A_ for the primary donor is far less than the value of ρ_A_ = 2.5 × 10^−3^ for the Chl_D1_ exciton in *chl-a*-PS II. A significant reduction in CD for the single FRL-chl in FRL-PS II can be expected regardless of where it is located. This is due to the relatively large energetic difference between the 5 *chl-a* pigments and the single FRL-chl in the reaction center (compared to the coupling matrix elements) reducing the ability of (unchanged) exciton coupling matrix elements to induce CD. The calculated value of ρ_A_ = −0.02 × 10^−3^ for the primary donor as Chl_D1_ may appear to be anomalously small given the significant exciton matrix elements between Chl_D1_ and P_D1_, P_D2_ and Pheo_D1_ (Table S4). This result highlights the fact that the CD of an exciton coupled multi-pigment array is not simply reducible to sums of individual couplings.

Preliminary calculations were done to explore the influence of proximal antenna-based FRL-chls on the CD in the 726 nm region. The closest pigment to Chl_D1_ present in CP43 is the ‘linker’ Chl508 [43]. Exciton coupling matrix elements between the reaction center pigments and Chl508 were calculated using a point-dipole approximation [41]. The largest matrix elements between Chl508 and the reaction center pigments were in the range of 1 cm^−1^ to 2 cm^−1^ and 2 orders of magnitude smaller than the largest matrix elements within the reaction center (Table S3). Exciton CD calculations were then performed for both the Chl_D1_ and P_D2_ options for the 726 nm pigment, whilst taking Chl508 to be the FRL-chl at 737 nm (Table S1). The CD calculated at 726 nm when the primary donor was Chl_D1_ became significant in magnitude but remained negative having ρ_A_= −0.32 × 10^−3^. With P_D2_ as the primary donor, the calculated CD including Chl508 remained positive with ρ_A_= 0.36 × 10^−3^ and close to the experimental value of 0.47×10^−3^. In both cases, the sign of the CD calculated for the 737 nm band was positive and thus not in agreement with experiment (see Fig. 6, Table S1 and Fig. S4).

In addition to exciton CD, there is the monomeric CD component of *chl* Q_y_ transitions. This is independent of, and additive to, exciton CD. Monomeric Q_y_ CD of *chl-a* is ρ_A_ −0.16 × 10^−3^ [47], −0.07 × 10^−3^ for *chl-d* [38] and −0.03 × 10^−3^ for *chl-f* [48]. These values are all small in magnitude and of the opposite sign to the observed value of ρ_A_ = 0.47 × 10^−3^ for the 726 nm band in FRL-PS II.

### 3.6 Arguments in favor of the primary donor being Chl_D1_

Chl_D1_ is considered to be the primary donor in *chl-a*-PS II [3, 5, 7, 44, 45]. Chl_D1_ is then the obvious pigment to exchange for a long wavelength pigment [1] to create a low energy primary donor. This apparently straightforward option is reinforced by the fact that all known FRL-PS II systems for which sequences are available show a pair of conserved amino acid changes D1Tyr120 and D1Thr155. These can be modelled as providing H-bonds to the extra formyl group that is the substituent on ring I of the chlorophyll that is responsible for the long-wavelength shift in a FRL-chl [1]. This seems to be a genetic/structural “smoking gun”. In contrast, amino acid residues facing the same position in the ring I of P_D2_ appear to be unable to form H-bonds having aliphatic side chains (Val and Leu). These residues were also found to be conserved in both the FRL and WL version of PSII. Nevertheless, we cannot exclude H-bond formation via the amide group of the peptide bond.

The mechanistic and redox arguments cited in the introduction are also strong. Given the similar rates and efficiency of O_2_ evolution, it is assumed that the donor-side components of FRL PS II are tuned to have the same redox potentials as in *chl-a*-PS II [1]. Thus, the primary electron donor FRL-chl^•+^/chl is taken as having a similar potential as its counterpart in *chl-a*-PSII. However, FRL-chl absorbing at 725nm will have ~110 meV less energy available in the excited state than does *chl-a* absorbing at ~680nm. This will result in the FRL-chl^•+^/chl*couple having higher potentials and therefore FRL-chl* will have less reducing power than its *chl-a* counterpart [1].

A less reducing FRL-chl* strongly favors pheo-a as the primary electron acceptor because the pheo-a/pheo-a^•-^ couple has a much higher reduction potential than that of *chl-a/chl-a^•-^.* The structure dictates that Pheo-a can only be the primary acceptor when Chl_D1_* is the primary donor and this provides a good argument for the primary donor being Chl_D1_ in FRL-PS II [1]. In models in which P_D2_ is the primary donor, the primary acceptor would be *chl-a* (either P_D1_ or Chl_D1_). This is expected to be unfavorable in redox terms. A similar argument was made for assigning the *chl-d* primary donor on Chl_D1_ in *Acaryochloris marina* PS II [4]. Along the same lines, recent calculations of the redox potentials of the chlorins in *chl-*a-PSII indicated that Chl_D1_* electron donation to Pheo is favorable, but that P_D1_* donation to Chl is energetically unfavorable [45]. If this trend is extrapolated to FRL-PSII, FRL-chl* is expected to be even less capable of reducing a *chl-a* primary acceptor.

The chemical argument mentioned in the introduction comes from the demonstration that chlorophyll has acid base properties that will favor ligation to water over histidine [46]. In the *chl-a*-PS II reaction center, only the Chl_D1_ and Chl_D2_ pigments have water coordination and are thus more likely as candidates for replacement by *chl-f* [1]. It is not possible to rule out histidine ligation to *chl-f*, in either the PS II reaction center or in peripheral antenna, but the chemical preference exists.

## 4. Conclusions

Illumination of FRL-PS II at 1.8 K at a series of wavelengths confirmed that the pigment at 726 nm is the primary electron donor. Charge separation occurred relatively efficiently at wavelengths ≤ 726 nm but became much less efficient at longer wavelengths. This verifies the conclusion made earlier [1].

A global analysis of absorption, CD and MCD data enabled the determination of the exciton spectral amplitudes, positions and widths of all 5 FRL-chl Q_y_ state spectral components in FRL-PS II. The Q_A_^•-^ induced electrochromism in FRL-PS II generated by low temperature and low fluence illumination was modelled by exciton coupling calculations (Figs. 5, 6 and S9). The starting point for these calculations was a new *quantitative* analysis of existing electrochromism data of *chl-a*-PS II core complexes prepared from spinach, allowing a model-free determination of the value of Δμ/f. The FRL-PS II electrochromic shift at 726 nm could be quantitatively accounted for by transferring the Chl_D1_ or P_D2_ *chl-a* site energy appropriate to spinach to the FRL 726 nm position, whilst leaving all exciton coupling terms, other site energies and geometric parameters as well as Δμ/f unchanged. The P_D2_ calculation provided a slightly better fit to the overall electrochromic pattern in the 650 nm to 690 nm region. The exciton modelling was also able to exclude the possibility that there was more than one FRL-chl in the reaction center, again in line with earlier ideas [1].

The CD calculated by the exciton modelling was able to account for the observed value of ρ_A_ =ΔA/A at 726 nm, without the use of additional parameters, when the primary donor was located at P_D2_. When located at Chl_D1_ the CD calculated was 30 times smaller and of the wrong sign in comparison to experiment. Preliminary exciton calculations of coupling between the reaction center chlorins and the closest chlorin in the proximal antennas indicate that coupling an FRL-chl at these locations is unlikely to account for the experimentally observed CD amplitude at 726 nm.

There are a number of arguments that make a good case for the Chl_D1_ being the location of the single FRL-chl in the reaction center. However, these arguments are intuitive, mechanistic and, when structural (i.e. the conserved amino acid changes) are yet to be verified by experiment. The current work presents initial experimental evidence, particularly from the concurrence of calculated and experimental CD at 726 nm, that the primary donor could be located at P_D2_.

Rather than speculating further on this surprising result, we look to perform additional spectroscopic experiments and calculations that may provide further evidence relevant to the character and location of the primary donor in FRL-PS II.

## Acknowledgements

We recognize the support of the Australian Research Council through grants DP110104565 and DP 150103137 (EK), FT140100834 (NC). This work was supported by BBSRC grants BB/L011506/1 and BB/R001383/1 (AWR, AF and DN).

## Supplementary Information

### SI-1 Allophycocyanin and FRL chlorophyll content of FRL-PS II

The quantification of the five far red chlorophylls in FRL-PS II presents a challenge. Fig. 2 of the manuscript identifies the presence of allophycocyanins which overlap the spectral region where the FRL chlorins, *chl-f* and *chl-d*, absorb.

Fig. S1 provides the integrals of absorption in the Q_y_ regions. Both high and low allophycocyanin samples have total absorption in the 700 nm - 800 nm region that exceeds a value corresponding to 5 chls by ~2-3 chl. The molar extinction of allophycocyanins isolated from *Synechocystis* is ~2×10^5^ M^−1^ cm^−1^ for both types of allophycocyanin found in this organism. Thus, there are likely to be ~1-2 allophycocyanin per FRL-PS II in our preparations.

**Fig. S1.**
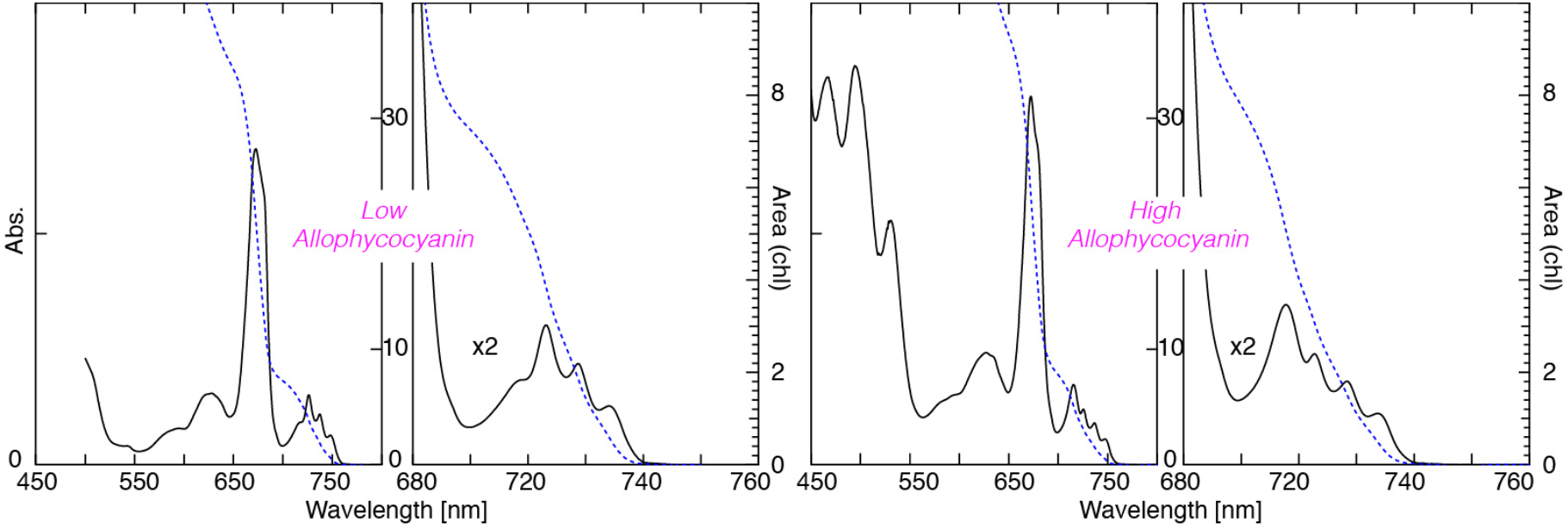
Absorption spectra of low allophycocyanin and high allophycocyanin FRL-PS II at 1.8 K along with their integrals (dashed curves), scaled to ~35 chl total Q_y_ absorbance.

Allophycocyanins are known to form both monomers and trimers [1] with trimers being red shifted. This variability precludes a quantitative assessment of the allophycocyanin components. Different monomer and trimer allophycocyanin content is likely to be responsible for the changing spectral profiles in absorption and CD of allophycocyanin components (see Fig. 3 of manuscript).

### SI-2 Fluorescence Excitation Spectra and Temperature Dependent Fluorescence

Fig. 1 of the manuscript highlights the difference between the emission spectra of the *chl-a*-PS II and that of FRL-PS II. In the latter the dominant peak is at 752 nm, with a weaker emission in the 720 nm region, attributable to allophycocyanin.

The spectra in Fig. S2 show that allophycocyanin absorption leads to a low level of emission from FRL-PS II. The excitation spectrum presented in Fig. S2 is similar to the MCD spectra of FRL-PS II in Fig. 1 of the manuscript in that the characteristic 714 nm allophycocyanin peak is missing. Absorption in the 680 nm to 710 nm region leads to the weaker emission peak at ~720 nm which does not correspond to the 714 nm absorption. This emission likely arises from a form of allophycocyanin (monomer, trimer etc.) present in the sample that is not functionally connected to FRL-PS II (see below).

**Fig. S2.**
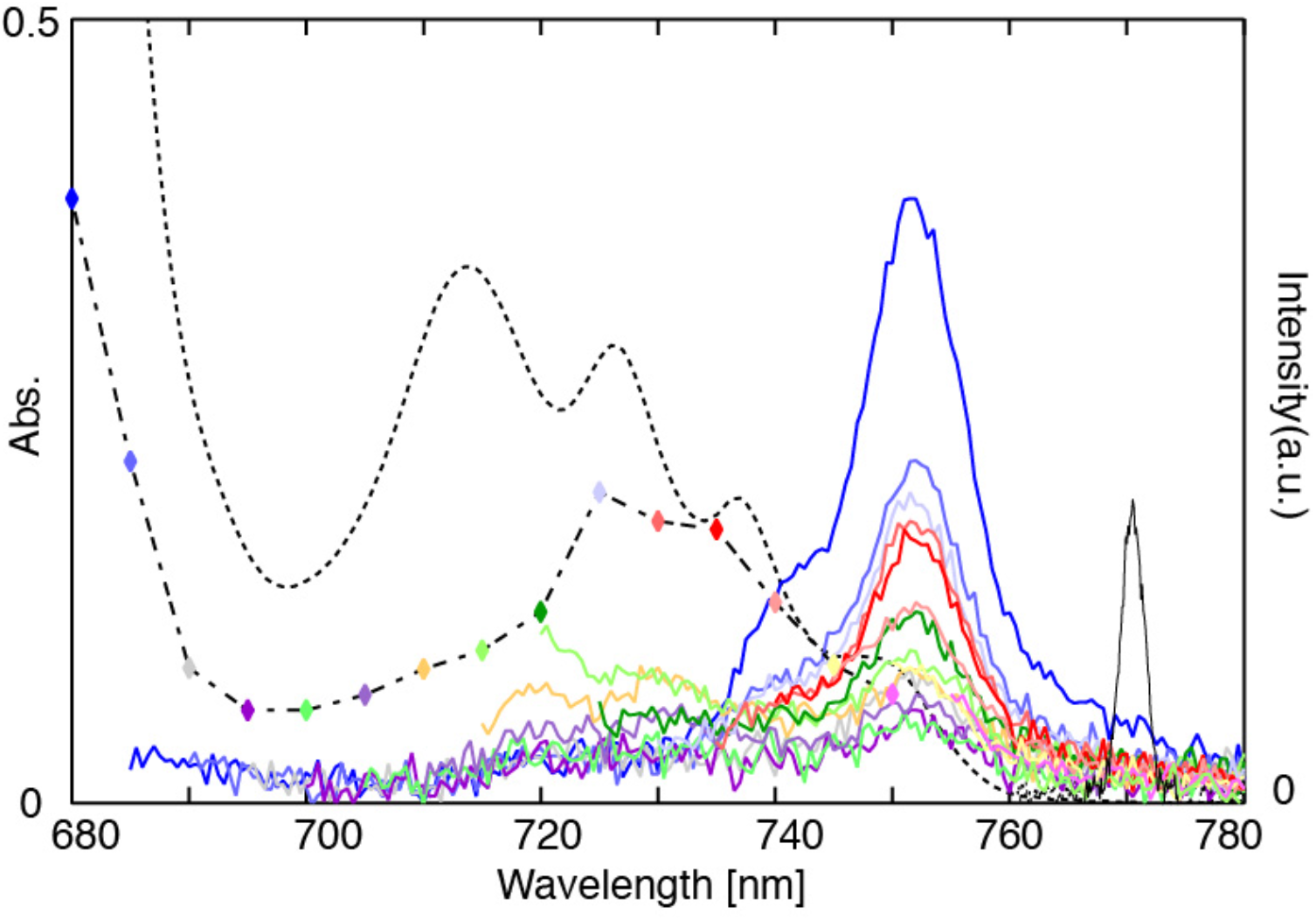
The absorption spectrum (dotted line) and the emission spectra (coloured lines) of FRL-PSII are shown. Spectra were taken at 77 K of a high-allophycocyanin FRL-PSII sample. The emission spectra were taken with excitation at different wavelengths in 5 nm steps from 680 nm to 755 nm. The corresponding excitation spectrum (composed of coloured diamonds which are joined by a double-dashed line) is generated by plotting the amplitude of the emission peak at 755 nm. The colour of each diamond in the excitation spectrum matches that of the corresponding emission spectrum. The narrow peak seen at 770 nm (black line) is obtained by measuring 770 nm exciting light. This spectrum serves to calibrate the excitation light bandwidth from the monochromator as ~2.4 nm FWHM.

Fig. S3 shows the excitation dependence of the well-resolved fluorescence in a high-allophycocyanin sample at 1.8 K. The weak emission from the contamination by *chl-a*-PS II in the 685nm – 693 nm region, and an allophycocyanin peak at 720 nm discussed above, are apparent, as well as the double peak feature at 741 nm and 752 nm associated with FRL-PS II. The relative intensities of the double peaks in FRL-PS II (Fig. S3) do not vary with excitation wavelength. This indicates that both peaks originate from the FRL PS II component of the FRL-PS II core complex. Additionally, the intensity of the 720 nm peak, when excited at different wavelengths, does not scale with that of the double peaks, indicating it arises from a allophycocyanin in the FRL-PS II preparation and a further indication that the allophycocyanin does not transfer all its excitation energy to a PS II core assembly.

**Fig. S3.**
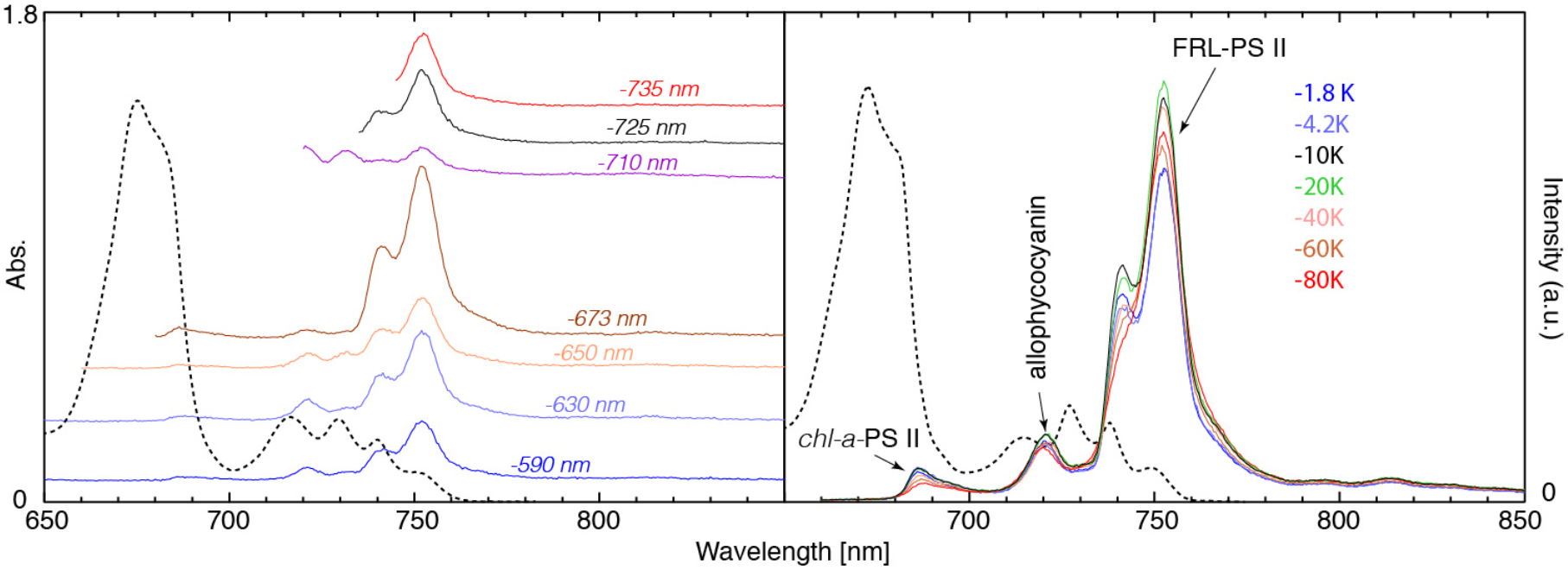
Excitation dependence of fluorescence from a high-allophycocyanin FRL-PS II preparation at 1.8 K (left panel). Temperature dependent emission of a medium-allophycocyanin sample excited at 532 nm (right panel). Absorption spectra (black dotted lines) are provided for comparison in both panels.

The temperature dependence of the high energy fluorescence component in the 680 nm to 700 nm region (Fig. S3) shows the temperature dependence of *chl-a*-PS II [2], which is clearly present as a contaminant in this sample. Assuming PS II core complexes of FRL-PS II and *chl-a*-PS II have comparable quantum efficiencies in emission and have equivalent absorption strengths at the excitation wavelengths used, the contamination level is estimated to be ~7 %. Fig. S3 identifies a modest decrease in the allophycocyanin fluorescence at 720 nm in the 1.8 K to 80 K range. It exhibits a simple broadening in spectral profile. This behavior is typical for an isolated antenna assembly such as CP43 [3] or CP47 [4].

The PS II core complex emission component of FRL-PSII, having emission peaks at 741 nm and 753 nm, shows a stronger and more complex temperature dependence. The two peaks have distinctly different temperature dependence, with the 742 nm peak reducing in amplitude with temperature more quickly than the 753 nm peak. This temperature dependent behavior is reminiscent of that shown by *chl-a*-PS II where the higher energy component can be attributed [2, 5, 6] to CP43 at the lowest temperatures. This peak becomes less prominent at higher temperatures. The barrier to excitation transfer to the reaction center is overcome by thermal activation at higher temperatures and consequently emission from the lowest energy excitations of CP43 becomes depleted.

### SI-3 Determination of the five FRL chlorophyll Q_y_ Bands

An initial estimation the peak positions and linewidths of the three distinct FRL chlorophyll features is straightforward. There is some indication of a high-energy shoulder near 713 nm in the MCD spectra (see Fig. 3 of the manuscript) and is also visible in the excitation spectrum shown in Fig. S2.

Attempts to fit MCD spectra with five Gaussians, one for each FRL chlorophyll, did not provide a uniquely defined fit. These attempts do indicate that two FRL Q_y_ absorptions have greater linewidths than the three obvious features. Absorption and CD spectra do not assist greatly in locating the positions and widths of the two remaining FRL chlorophyll molecules, owing to interference by allophycocyanins.

The amplitude of the MCD relative to the absorption (B/D) of chlorins is known to vary significantly [2, 7–10] with the primary donor having a greatly reduced value of B/D. The approach developed was to subtract a scaled absorption spectrum from the corresponding MCD spectrum and vary the scaling factor until each well-resolved FRL chlorophyll feature in the MCD spectrum was nulled. This procedure worked well and provided relative B/D ratios for each of the three distinct FRL chlorophyll Q_y_ absorptions.

These pre-determined relative B/D ratios were then held constant in a five band Gaussian fitting of the MCD spectrum. This process determined the position, width and relative B/D of the two remaining FRL chlorophylls, i.e. Chl-3 and Chl-5, as shown in Table S1. The same positions, widths and relative B/D values were obtained in independent analyses of both high and low allophycocyanin MCD spectra.

**Table S1.**
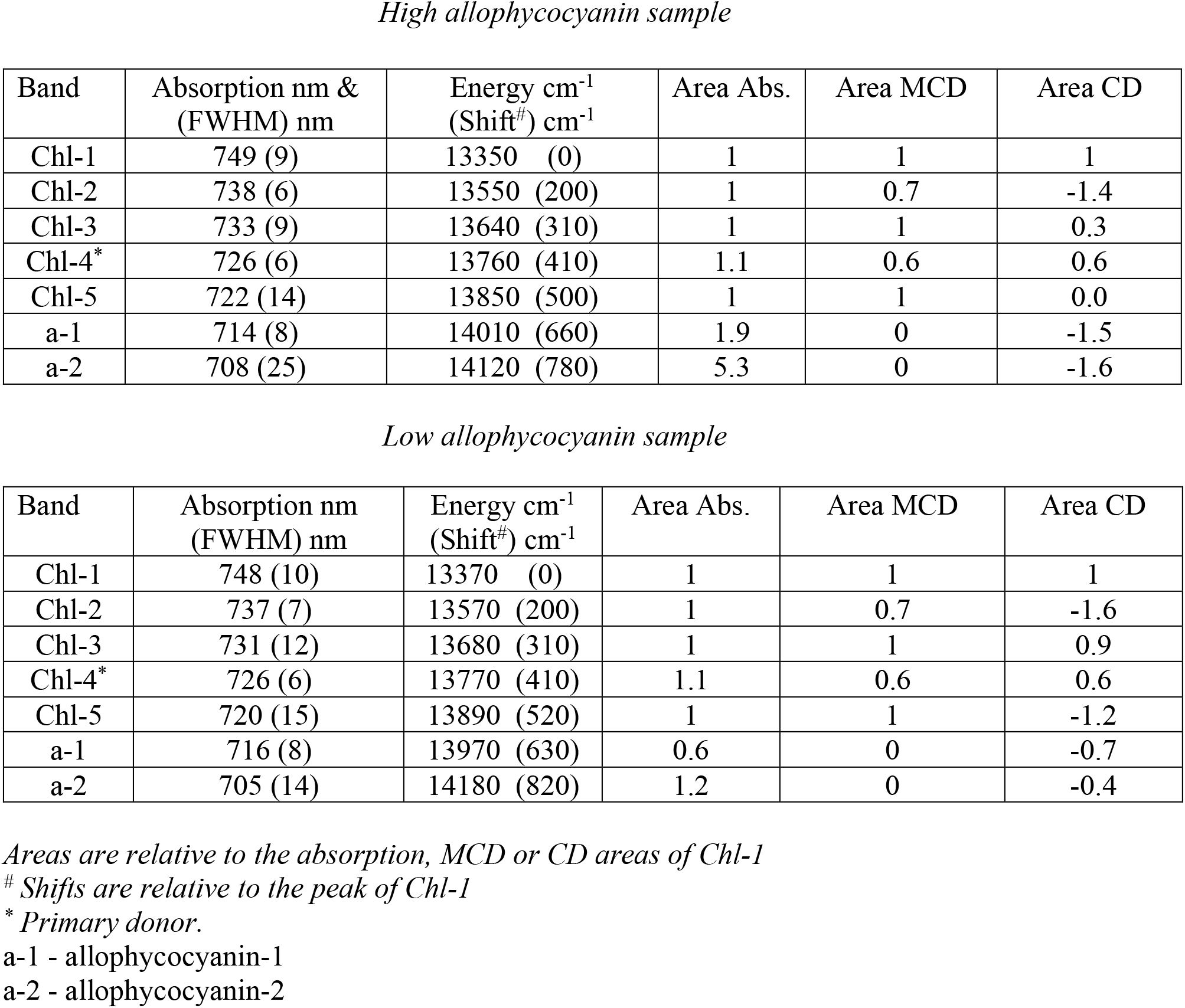
FRL-PS II Global Analysis

The positions and widths of Chl-3 and Chl-5 were not dependent on the precise value of their relative B/D values (these are proportional to the ‘Relative Area MCD’ column in Table S1). The best overall fit was obtained when Chl-1, Chl-3 and Chl-5 had equal B/D values. The primary donor (Chl-4) had a smaller B/D value. This is consistent with previous observations of B/D for the primary donor in *chl-a*-PS II [2, 7–10] although the full chemical significance of this notable electronic characteristic of primary donors is not understood.

Having determined the position and width of each FRL chlorophyll Q_y_ excitation (Table S1), the FRL chlorophyll Q_y_ absorption spectrum was then fitted to five bands of equal area. This process involved a single linear fitting parameter. Two Gaussians were introduced to account for the allophycocyanins absorptions (allophycocyanin-1 and allophycocyanin-2 in Table S1). The narrow allophycocyanin component (allophycocyanin-1) in high-allophycocyanin and low-allophycocyanin samples are similar but allophycocyanin-2 is quite different. The fits for allophycocycanin-1 and allophycocyanin-2 -shown in Fig 3 of the manuscript and Table S1 remain indicative only, as there is significant correlation between fitting parameters. The presence of different forms of allophycocyanins was discussed in SI-1.

In Table S1 the relative area of the Chl-4 absorption was revised to have a value of 1.1 chl. Exciton modelling (SI-4) establishes that the interactions between the FRL chlorophyll primary donor and its *chl-a* neighbors within the reaction center leads to this (relatively low) level of transfer of intensity.

The CD spectra were then fitted, with the amplitude of each Q_y_ CD feature allowed to vary. The allophycocyanin CD components were allowed unconstrained amplitudes, widths and positions. The widths and positions of allophycocyanin CD features are not those as determined by the corresponding fit of the absorption spectrum. For the five FRL chlorophyll Q_y_ excitations, each is an individual exciton Q_y_ component. The CD profile of a single purely electronic Q_y_ excitation follows that seen in absorption [11]. The absorption profiles of allophycocyanins are composite of a number of exciton-coupled pigment excitations. Thus allophycocyanin CD profiles do not follow those in absorption [1, 12].

The right-hand columns in Table S1 show that the area of Q_y_ CD features associated with Chl-1, Chl-2 and Chl-4 reproduce well between the high and low allophycocyanin samples, whereas discrepancies appear for Chl-3 and Chl-5. Fig S4 provides a comparison of the experimental and fitted CD for high and low allophycocyanin samples, along with the individual CD components determined by the fitting procedure described above.

**Fig. S4.**
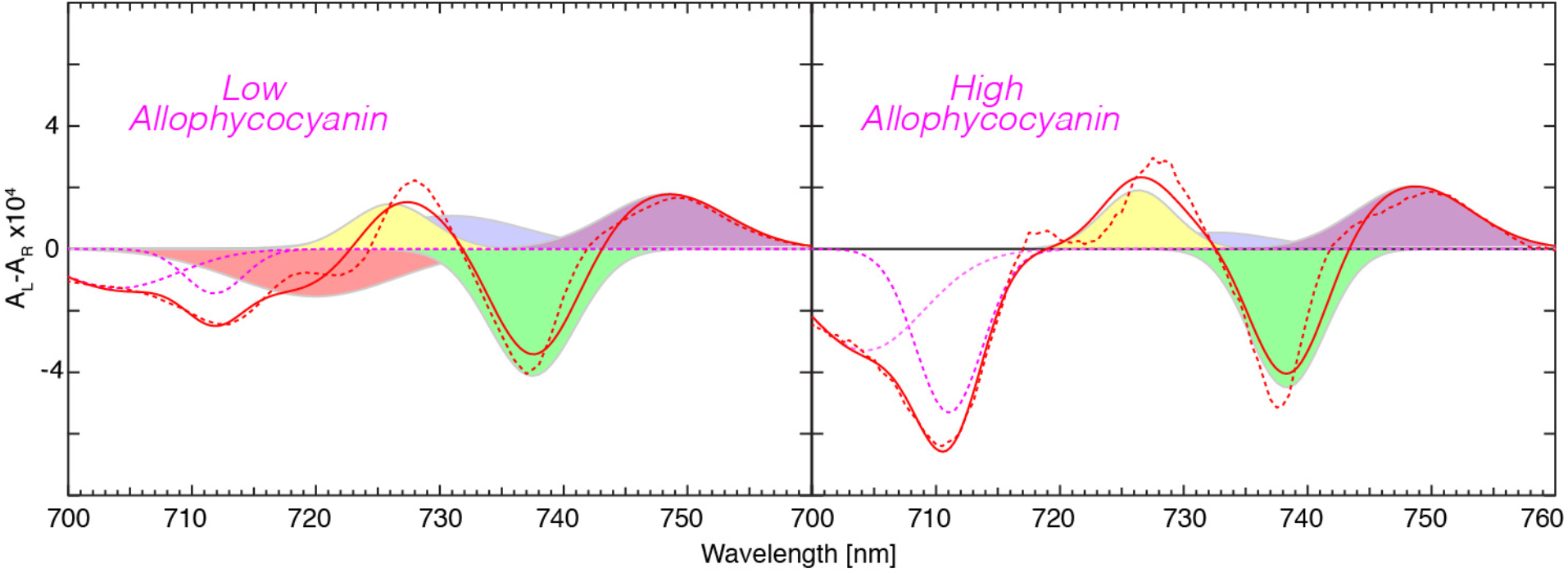
The 1.8 K experimental CD and corresponding fitted CD spectra of low and high allophycocyanin FRL-PS II samples are shown as dotted and solid red lines respectively. These spectra are also presented in Fig. 3 of the manuscript in less detail and are displayed in conjunction with corresponding absorption and MCD data and analysis. The fitted CD components (see text) are displayed as coloured overlapping Gaussians using the same colour scheme as used in Fig. 3.

A significant discrepancy is that Chl-5 is attributed to have minimal CD in the high allophycocyanin sample but a strong negative CD in the low allophycocyanin sample. By contrast, the three sharp FRL-chl features, Chl-1, Chl-2 and Chl-4 are well determined in both samples and their CD amplitudes are not significantly correlated. However, the fit of Chl-3, Chl-5, a-1 and a-2 CD features are significantly correlated, particularly for the low allophycocyanin CD data. The region of major discrepancy (720 nm) shows a sharp inflection in both samples that is not reproduced in either absorption or MCD data. A more complete representation of allophycocyanin CD thus requires extra components in the fitting procedure. Additionally, rather than a simple reduction of allophycocyanin content between the two samples, it seems that extra (allophycocyanin) assemblies are introduced via the procedure used to reduce the total allophycocyanin content. These assemblies appear to have significant negative CD in the 720 nm to 730 nm region, making it impossible to disassociate it from CD associated with Chl-5. Without further experimental information, we cannot determine the CD of Chl-5. It seems likely that the CD of Chl-3 is significant and positive.

### SI-4 Electrochromic Shifts

A comparison of Q_A_^•-^ formation-induced electrochromic shift patterns for a range of organisms compared to those seen in FRL-PS II, is provided in Fig S5.

**Fig. S5.**
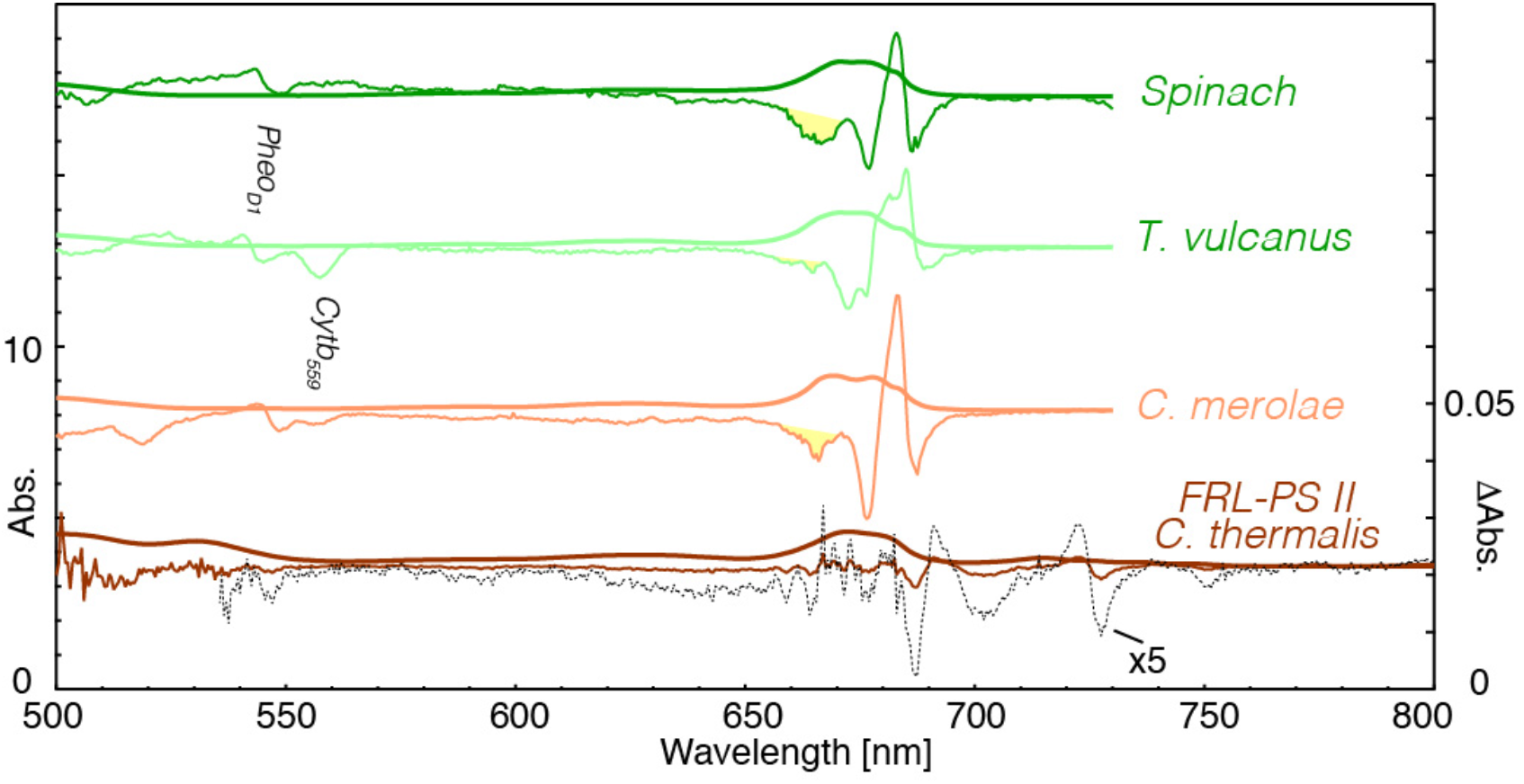
Absorption (left Scale) and illumination-induced difference spectra (right scale) for PS II core complexes derived from Spinach, T. vulcanus [13], C. merolae [14] and FRL C. thermalis [15] at 77 K. The shaded yellow regions are assigned to ChlZ_D2_ bleaching (see text). Green light illumination fluences were standardized at 300 s of 5 mW/cm^2^. Spectra have been scaled to comparable Q_y_ absorption intensities and offset for viewing. Q_A_^•-^ yields in the first three systems are similar (60-80%), whereas the corresponding yield in FRL-PS II is ~10 to 20 times less.

Low temperature electrochromic shifts associated with Q_A_^•-^ formation have been extensively studied [16–21]. In Fig. S5, similar Q_x_ ‘C550’ electrochromic blue-shifts [16] of Pheo_D1_ are evident in all organisms. The nearby bleach feature associated with oxidation of reduced *cytb_559_* is evident in *T. vulcanus* and *C. merolae* preparations. The FRL-PS II preparation of *C. thermalis* used for Fig. S5 did not contain a significant fraction of reduced *cytb_559_* and thus no bleach is seen, although this spectral feature was evident at 556.5 nm in other sample preparations (see Fig. S6 and [15]). Fig. S5 also shows the bleach near 666 nm, highlighted as a shaded yellow region, associated with the oxidation of a *chl-a* (often considered to be ChlZ_D2_) for the first three organisms. This oxidation process appears less pronounced in FRL-PS II and other secondary donors such as β-carotenes may be involved [22–24].

**Fig. S6.**
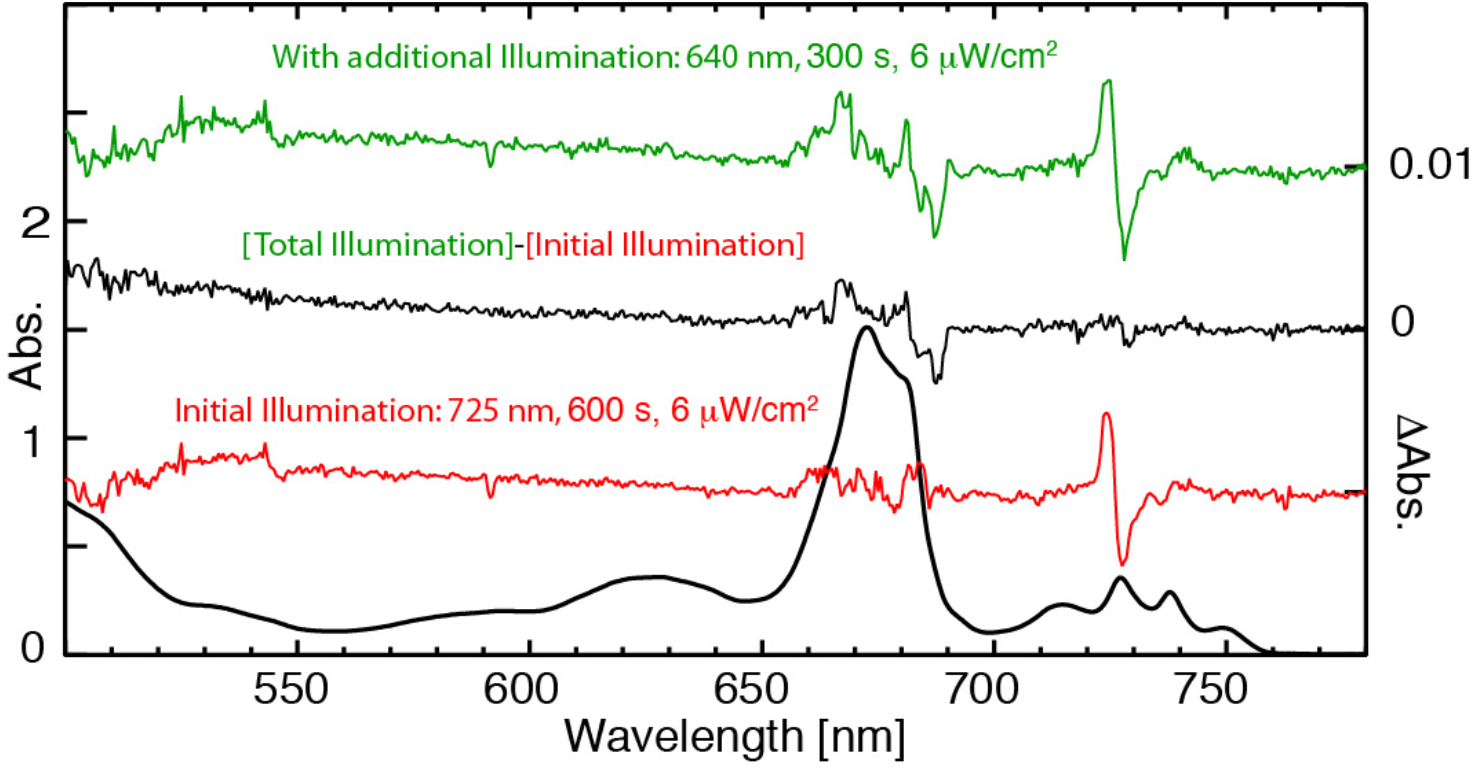
The bottom black trace is the absorption spectrum (left. scale) of the low-allophycocyanin FRL PS II core complex at 1.8 K. The red trace is the electrochromism spectrum (right scale) induced by illumination at 725 nm with the fluence indicated. The green trace is the accumulated electrochromism after an additional illumination at 640 nm. The thin black trace is the change in electrochromism induced by the 640 nm illumination component.

The dominant electrochromic features in the Q_y_ region for *chl-a*-PS II have been identified as a red shift of Pheo_D1_ near 680 nm and blue shift of Chl_D1_ Q_y_ near 685 nm. The precise position and shape of shift features vary with organism. This variation is due to both small changes in pigment site energies with organism, along with changes in the effects of excitonic interactions between pigments associated with relatively small changes in their site energies. This is because the relative energy of sites can be changed quite significantly with respect to the exciton coupling matrix element between pigments. Shift features are doubled in some preparations of *T. vulcanus and T. elongatus i.e.* when there are two forms of the D1 protein expressed in the sample [25].

In the FRL-PS II Q_y_ region, the largest shifted feature is well separated with an intense blue shift feature occurring at 726 nm [15]. Compared to the data in Fig. S5, which are taken at 77 K, those taken at 1.8 K (Fig. 2 and Fig. 4 of manuscript) show better-resolved shift features.

FRL-PS II core samples contain some *chl-a*-PS II, as evidenced by its characteristic emission spectrum appearing in Fig. 1 of the main manuscript. FRL-PS II samples illuminated with light at wavelengths significantly shorter than 700 nm, such as the green light used to generate difference spectra in Fig. S5, can be expected to display electrochromism associated with the *chl-a*-PS II component, as well as that of the (majority) FRL PS II component. Illumination of *chl-a*-PS II at 725 nm and 1.8 K is known to induce Q_A_^•-^ formation (and as evidenced by electrochromism) but with low quantum efficiency [20, 26]. The *chl-a*-PSII electrochromism induced by illumination in the 700 nm to 730 nm range is due to absorption by the weak and broad, deep-red charge transfer state of *chl-a*-PS II [14, 20, 26].

Fig. S6 demonstrates that illumination of a low allophycocyanin FRL-PS II core complex sample with 640 nm light, following initial illumination at 725 nm, induces minimal additional electrochromism at 726 nm. The light-induced Q_A_^•-^ formation of the FRL PS II component was essentially completed by the first illumination. However, additional 640 nm illumination leads to an electrochromism spectrum with a negative feature near 686 nm and a positive component near 682 nm. The spectrum exhibits low signal to noise as it was generated as a double-difference spectrum and also recorded in a region of high and rapidly varying absorbance. However, the pattern is compatible with it being due to the Chl_D1_ blue-shift of *chl-a*-PS II *of C. thermalis* [15]. For comparison, the Q_A_^•-^ induced electrochromic shift of *T. vulcanus* (Fig. S5), a *chl-a* PS II thermophilic cyanobacterium has negative and positive peaks at 688 nm and 684 nm respectively.

The red trace in Fig. S6 shows no evidence of the negative feature seen near 685 nm in the black trace. We can conclude that no significant component of the *chl-a*-PS II contaminant of the FRL-PS II sample is photoconverted by 725 nm light. The electrochromism pattern in the red trace is then due only to the FRL PS II majority component in the sample.

The linewidth of the 726 nm feature (7 nm or ~130 cm^−1^ FWHM) allows an initial estimate of the electrochromic shift to be determined, independent of knowledge of the Q_A_^•-^ yield. By measuring the offset, δ, of the derivative of the (initial) absorption spectrum compared to the electrochromic shift feature, the electrochromic shift would then be 2δ, i.e. 2.2 nm or equivalently 42 cm^−1^. This assumes that the relatively small fraction of FRL PS II that undergoes photoconversion has the same spectral distribution as the majority. Those FRL-PS II centers that are not photo-converted to the trapped Q_A_^•-^ form do not contribute to the shift feature. The shift feature has a crossing point independent of yield as seen in Fig. S7, and is displaced 1.1 nm from the peak of the absorption, taken before illumination.

**Fig. S7.**
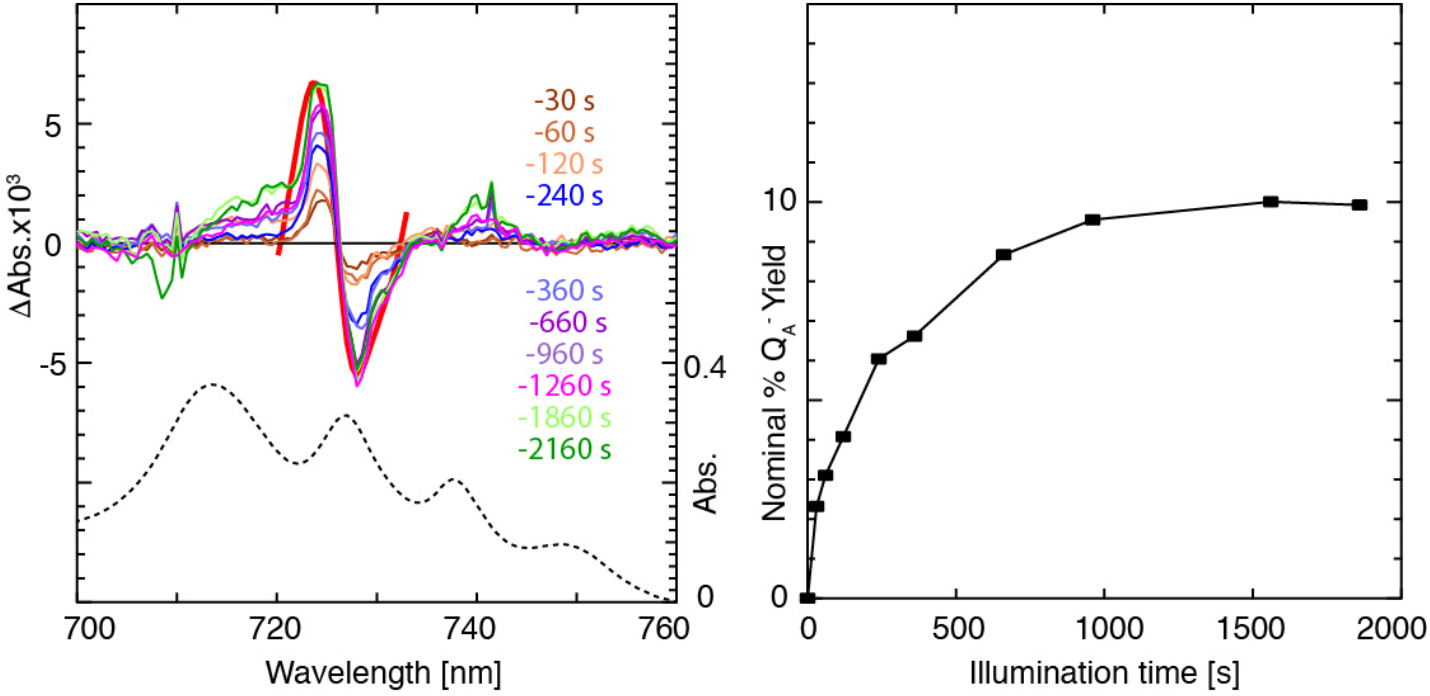
The left panel shows the fluence dependence of the electrochromic shift of high-allophycocyanin FRL-PS II at 1.8 K illuminated with 6 μW/cm^2^ power density light at 725 nm. The red trace in the 726 nm region is the synthetic shift feature created by displacing the initial absorption spectrum by 2.2 nm (42 cm^−1^) to the blue and subtracting this from the original spectrum. The right panel is the nominal yield of photoconversion to the Q_A_^•-^ form of the enzyme versus illumination time, assuming a 2.2 nm electrochromic shift (see revision in text).

**Fig. S8.**
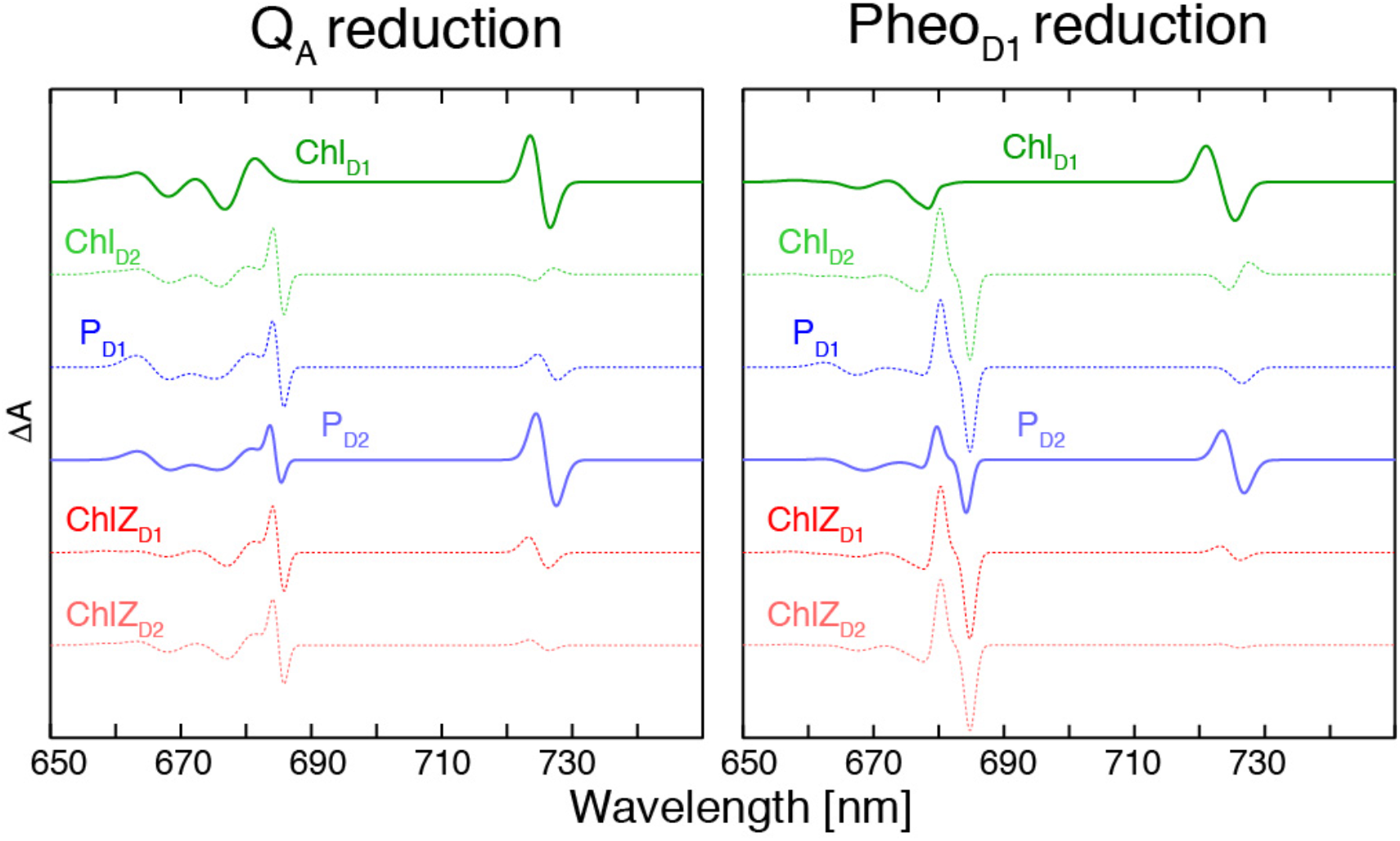
Left and right panels show calculated electrochromic shift patterns induced by an electron located at the Q_A_ and Pheo_D1_ positions, respectively. Shift patterns were calculated using the parameters in Tables S2 and S3. Calculations are presented for all locations of a single FRL chlorophyll in the reaction center. The two cases which induce an electrochromic shift near 726 nm compatible with experimental data (Chl_D1_ and P_D2_) are accentuated (thicker solid lines).

A Q_A_^•-^ yield can then be estimated by simulating a 100% yield shift feature at 726 nm. This artificial, full-yield spectrum is generated by shifting the absorption spectrum taken before illumination to the blue by 2.2 nm. Subtracting the shifted spectrum from the initial spectrum yields a spectrum corresponding to the entire 726 nm feature being affected. Scaling this shift feature against the experimental 726 nm shift amplitude then provides a Q_A_^•-^ yield. Fig. S7 shows fluence-dependent shift spectra with the apparent yield saturating at 10% when using 6 μW/cm^2^ excitation at 725 nm.

At the lowest yields, a simple 726 nm shift feature dominates. It is narrower than the synthetic shift feature. At higher fluences, shoulders develop which may be associated with photoconversion of a population of FRL-PS II that exhibits the full primary donor spectral distribution. The initial slope of the conversion curve in Fig S7 can be analyzed by gauging the number of absorbed photons for a given excitation fluence. This leads to a direct estimate of the initial quantum efficiency of the Q_A_^•-^ conversion as being close to unity. Such high initial quantum efficiencies were also seen in previous studies on spinach PS II core complexes [20] by monitoring the amplitude of the Q_x_ shift of Pheo_D1_.

However, at 77 K, shift features at both high and low fluence are displaced from the peak maximum by 0.5 nm, whilst a corresponding synthetic shift feature generated by a 0.5 nm displacement of the 77 K 726 nm absorption accurately reproduces the shift feature shape. This revised estimate of the shift of 0.5 nm leads to a corresponding 2 x increase in our estimate of the Q_A_^-^ yield (i.e. to 20%), compared to that proposed in Fig. S7.

As mentioned above, the low fluence shift feature at 1.8 K arises from a narrower component of the overall primary donor inhomogeneous absorption profile. This narrower component is displaced by 1.1 nm from the peak maximum. This displacement, combined with an actual electrochromic shift of 1 nm, leads to the observed 1.8 K value of 2δ of 2.2 nm. The narrower component has a linewidth close to 4 nm, rather than the overall linewidth of 7-8 nm as given in Table S1. This narrower component of Chl-4 is more in line with the width observed for the Chl_D1_ site energy linewidth in the *chl-a*-PS II core complex (from spinach), which is 1.6 nm (Table S2).

**Table S2.** Spinach Core Complex PS II reaction center site parameters

|  | Site Energy nm/cm<sup>-1</sup> | FWHM Linewidth nm/cm<sup>-1</sup> |
| --- | --- | --- |
| Chl<sub>D1</sub> | 684/14620 | 1.6/40 |
| Pheo<sub>D2</sub> | 678/14750 | 4.0/100 |
| Pheo<sub>D1</sub> | 676.5/14780 | 5.6/140 |
| P<sub>D1</sub> | 668.5/14960 | 6.3/140 |
| P<sub>D2</sub> | 668.5/14960 | 6.3/140 |
| Chl<sub>D2</sub> | 669/14950 | 6.1/140 |
| ChlZ<sub>D1</sub> | 672/14880 | 6.1/140 |
| ChlZ<sub>D2</sub> | 665/15040 | 6.1/140 |

### SI-5 Exciton Modelling

Here, we used the same methodology previously used to model the PS II reaction center as present in core complexes of a number of organisms [27] and also for *isolated* PS II (RC-6 and RC-5) [10, 28] reaction centers. We calculated absorption and CD spectra of a range of possible configurations of FRL-PS II. Apart from the single FRL chl site, the reaction center parameters used were those for *chl-a*-PS II core complexes isolated from spinach as shown in Table S2. The coupling matrix used is provided in Table S3.

**Table S3.** PS II reaction center exciton coupling matrix elements [29]

| Coupling (cm<sup>-1</sup>) | Chl<sub>D1</sub> | Pheo<sub>D2</sub> | Pheo<sub>D1</sub> | P<sub>D1</sub> | P<sub>D2</sub> | Chl<sub>D2</sub> | ChlZ<sub>D1</sub> | ChlZ<sub>D2</sub> |
| --- | --- | --- | --- | --- | --- | --- | --- | --- |
| Chl<sub>D1</sub> | 0 | -2.18 | 43.51 | -27.32 | -46.77 | 3.54 | 1.67 | -0.09 |
| Pheo<sub>D2</sub> | -2.18 | 0 | 1.55 | 12.61 | -2.99 | 41.65 | -0.19 | -2.57 |
| Pheo<sub>D1</sub> | 43.51 | 1.55 | 0 | -3.96 | 15.06 | -2.37 | -2.52 | -0.18 |
| P<sub>D1</sub> | -27.32 | 12.61 | -3.96 | 0 | 158 | -41.83 | 0.45 | 0.58 |
| P<sub>D2</sub> | -46.77 | -2.99 | 15.06 | 158 | 0 | -22.04 | 0.6 | 0.61 |
| Chl<sub>D2</sub> | 3.54 | 41.65 | -2.37 | -41.83 | -22.04 | 0 | -0.05 | 1.8 |
| ChlZ<sub>D1</sub> | 1.67 | -0.19 | -2.52 | 0.45 | 0.6 | -0.05 | 0 | 0.15 |
| ChlZ<sub>D2</sub> | -0.09 | -2.57 | -0.18 | 0.58 | 0.61 | 1.8 | 0.15 | 0 |

Transition dipole strengths were taken from reference [30] as 5.47 D for *chl-a* and 4.25 D for *pheo-a* and structural data determined from crystal structure data [31].

Electrochromic shifts, Δ, induced by reducing Q_A_ and Pheo_D1_ forming their anion radical states, can be modelled by considering the interaction of 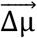 of each reaction center pigment with the electric field associated with a point charge at the position of the reduced species.

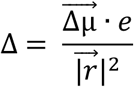

A starting point [30, 32, 33] is to assign Δμ to be 1.8 D for all *chl-a* Q_y_ excitations and 1.35 D for *pheo-a* Q_y_ excitations, with *e* being the electronic charge. This shift, Δ, is then reduced to Δ/f because of the influence of the local dielectric in the protein. When *f* is taken as 3, the site energy shifts are as given below.

**Table S4.** PS II Reaction Center Site Energy Shifts for f = 3

| Shift/cm<sup>-1</sup> | $Q_A^{\bullet-}$ | $Pheo_{D1}^{\bullet-}$ |
| --- | --- | --- |
| Chl<sub>D1</sub> | 14.4 | 77.2 |
| Pheo<sub>D2</sub> | -9.6 | -0.1 |
| Pheo<sub>D1</sub> | -53.8 |  |
| P<sub>D1</sub> | 5.8 | 0.4 |
| P<sub>D2</sub> | 18.0 | 45.1 |
| Chl<sub>D2</sub> | -5.4 | -18.8 |
| ChlZ<sub>D1</sub> | 10.0 | 11.2 |
| ChlZ<sub>D2</sub> | 3.9 | 2.8 |

The calculated electrochromic patterns shown in Fig. S8 indicate that Pheo_D1_ reduction experiments could assist in discriminating between the Chl_D1_ and P_D2_ locations for the primary donor. A relatively large primary donor shift for the Chl_D1_ assignment is calculated for the Pheo_D1_ reduction compared to that for the Q_A_^•-^ reduction. More useful may be that the associated electrochromic pattern seen in the *chl-a* Q_y_ spectral region is different for the Chl_D1_ and P_D2_ primary donor assignments. The room temperature Pheo_D1_ reduction spectra previously published in Fig 2D of [15] display a broad negative feature at 658 nm as well a stronger broad positive feature near 650 nm. Neither of these features correspond to the calculated electrochromism (see Fig. S8). In experiments [27, 35] on canonical *chl-a*-PS II preparations, there is no evidence of broad features similar to those seen in Fig. 2D of ref [15]. It thus seems important to generate the trapped Pheo^−^ state at low temperature in future experiments.

The signal-to-noise ratio and other limitations in the in the 725 nm-induced electrochromism spectra presented in Figs, 2 and 4 of the manuscript and in Fig. S9 makes it difficult to determine which calculated Q_A_ reduction pattern calculated (Fig. S8) better fits the experimental data in the *chl-a* region. Fig. S9 addresses this difficulty by adding results from identical experiments performed on three different FRL-PS II samples exhibiting differing allophycocyanin concentrations. Illuminations were performed with an identical fluence of 6 μW/cm^2^ at a wavelength of 725 nm for 300s. This illumination protocol is shown to correspond to conditions where no significant *chl-a* based photochemistry is induced (see Fig. S6), i.e. that associated with *chl-a*-PS II present in the samples. A comparison of the averaged shift pattern with those calculated for the 680 nm region appears to be more compatible with P_D2_ being the primary donor. The main feature at ~ 680 nm appears experimentally as a blue shift, as calculated for the P_D2_ case, rather than a red shift as calculated for the Chl_D1_ case.

**Fig. S9.**
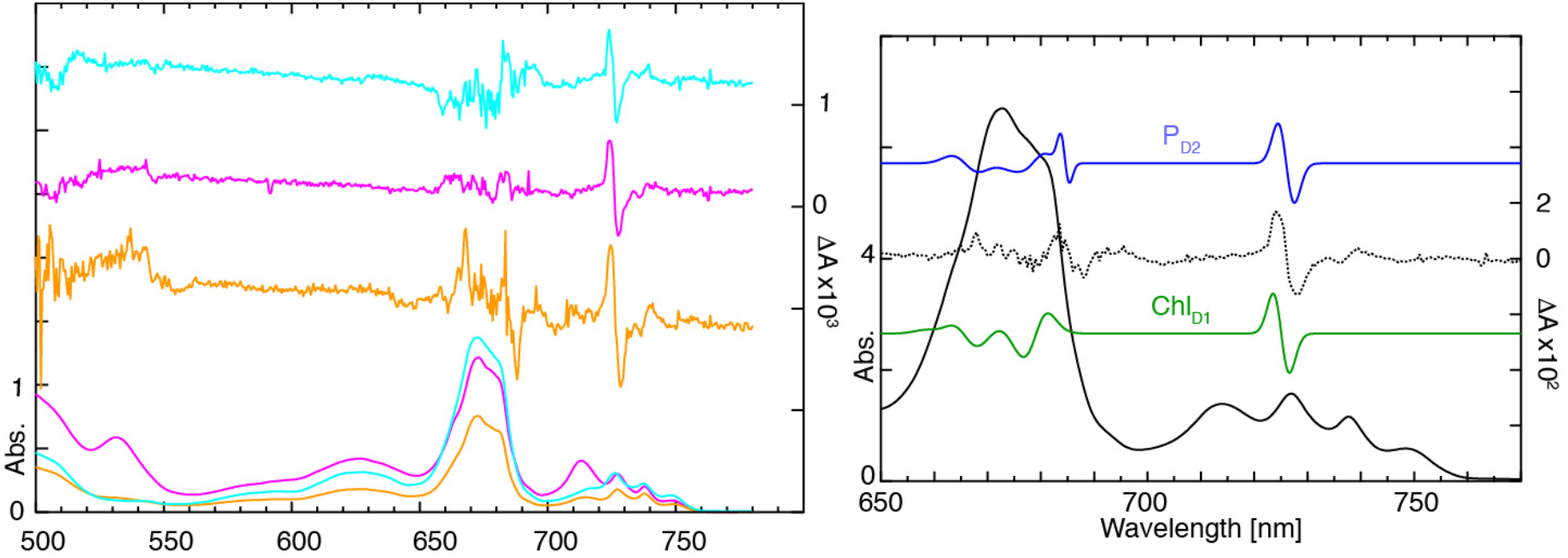
The left panel shows the absorption (lower traces) and electrochromic shift patterns (each offset for clarity) induced by 725 nm illumination at 1. 8 K of three different FRL PS II core complex samples having various allophycocyanin content. The right panel shows the sum of the three absorption spectra (thick black trace) and the sum of the corresponding three electrochromism spectra (thin black trace). Also shown are the calculated electrochromic patterns (from Fig. S8) when the primary donor is located at Chl_D1_ (green trace) and P_D2_ (blue trace). For both cases the calculated patterns have been scaled to match the experimental electrochromism amplitude.

## Abbreviations

chl-a-PS II: Canonical chlorophyll-a based Photosystem II
FRL-PS II: Far-red light grown PS II
CD: Circular Dichroism
MCD: Magnetic Circular Dichroism
*chl-a*: chlorophyll-a
*chl-d*: chlorophyll-d
*chl-f*: chlorophyll-f
FRL-chl: Far-red light chlorophyll (chlorophyll-d and/or chlorophyll-f)
SI: Supplementary Information

